# CD8^+^ T-Cell Metabolic Rewiring Defined by Single-Cell RNA-Sequencing Identifies a Critical Role of ASNS Expression Dynamics in T-Cell Differentiation

**DOI:** 10.1101/2021.07.27.453976

**Authors:** Juan Fernández-García, Fabien Franco, Sweta Parik, Antonino A. Pane, Dorien Broekaert, Joke van Elsen, Ines Vermeire, Thomas van Brussel, Rogier Schepers, Elodie Modave, Tobias K. Karakach, Peter Carmeliet, Diether Lambrechts, Ping-Chih Ho, Sarah-Maria Fendt

**Affiliations:** Laboratory of Cellular Metabolism and Metabolic Regulation, VIB–KU Leuven Center for Cancer Biology, VIB, Leuven, Belgium; Laboratory of Cellular Metabolism and Metabolic Regulation, Department of Oncology, KU Leuven and Leuven Cancer Institute (LKI), Leuven, Belgium; Department of Oncology, University of Lausanne, Lausanne, Switzerland; Ludwig Institute of Cancer Research, University of Lausanne, Lausanne, Switzerland; Laboratory of Myeloid Cell Immunology, VIB Center for Inflammation Research, Brussels, Belgium; Laboratory of Cellular and Molecular Immunology, Vrije Universiteit Brussel, Brussels, Belgium; Laboratory for Translational Genetics, VIB–KU Leuven Center for Cancer Biology, VIB, Leuven, Belgium; Laboratory for Translational Genetics, Department of Human Genetics, KU Leuven, Leuven, Belgium; Laboratory of Angiogenesis and Vascular Metabolism, VIB–KU Leuven Center for Cancer Biology, VIB, Leuven, Belgium; Laboratory of Angiogenesis and Vascular Metabolism, Department of Oncology, KU Leuven and Leuven Cancer Institute (LKI), Leuven, Belgium; Laboratory of Angiogenesis and Vascular Heterogeneity, Department of Biomedicine, Aarhus University, Aarhus, Denmark; State Key Laboratory of Ophthalmology, Zhongshan Ophthalmic Center, Sun Yat-Sen University, Guangzhou, Guangdong, P.R. China; Janssen Pharmaceuticals, Beerse, Belgium; Laboratory for Translational Research in Gastrointestinal Disorders, Department of Chronic Diseases and Metabolism, KU Leuven, Leuven, Belgium; Department of Pharmacology, Faculty of Medicine, Dalhousie University, Halifax, NS, Canada

## Abstract

Cytotoxic T cells dynamically rewire their metabolism during the course of an immune response. While T-cell metabolism has been extensively studied at phenotypic endpoints of activation and differentiation, the underlying dynamics remain largely elusive. Here, we leverage on single-cell RNA-sequencing (scRNA-seq) measurements of *in vitro* activated and differentiated CD8^+^ T cells cultured in physiological media to resolve these metabolic dynamics. We find that our scRNA-seq analysis identifies most metabolic changes previously defined in *in vivo* experiments, such as a rewiring from an oxidative to an anabolism-promoting metabolic program during activation to an effector state, which is later reverted upon memory polarization. Importantly, our scRNA-seq data further provide a dynamic description of these changes. In this sense, our data predict a differential time-dependent reliance of CD8^+^ T cells on the synthesis *versus* uptake of various non-essential amino acids during T-cell activation, which we corroborate with additional functional *in vitro* experiments. We further exploit our scRNA-seq data to identify metabolic genes that could potentially dictate the outcome of T-cell differentiation, by ranking them based on their expression dynamics. Among the highest-ranked hits, we find asparagine synthetase (*Asns*), whose expression sharply peaks for effector CD8^+^ T cells and further decays towards memory polarization. We then confirm that these *in vitro Asns* expression dynamics are representative of an *in vivo* situation in a mouse model of viral infection. Moreover, we find that disrupting these expression dynamics *in vitro*, by depleting asparagine from the culture media, delays central-memory polarization. Accordingly, we find that preventing the decay of ASNS by stable overexpression at the protein level *in vivo* leads to a significant increase in effector CD8^+^ T-cell expansion, and a concomitant decrease in central-memory formation, in a mouse model of viral infection. This shows that ASNS expression dynamics dictate the fate of CD8^+^ T-cell differentiation. In conclusion, we provide a resource of dynamic expression changes during CD8^+^ T-cell activation and differentiation that is expected to increase our understanding of the dynamic metabolic requirements of T cells progressing along the immune response cascade.

## INTRODUCTION

The adaptive immune system, and in particular cytotoxic (CD8^+^) T cells, confer an organism with short- and long-term protection against both foreign and intrinsic threats, such as viruses or tumors (Parham, 2014). Upon the onset of an immune response, resting naïve T cells in secondary lymphoid organs will become activated, undergoing a rapid proliferation burst and differentiating into an effector state (Cui and Kaech, 2010). Effector T cells will then migrate to the infection site, and deploy their cytotoxic activity against the infected or transformed cells. Following clearance, the effector population will further undergo a strong contraction, giving rise to a small population of long-lived memory T cells (Obar and Lefrançois, 2010). The latter will be able to readily re-activate in response to further antigenic stimuli, eliciting a faster and stronger secondary response against previously encountered pathogens or relapsing tumors (Veiga-Fernandes et al., 2000). The major phenotypical changes displayed by CD8^+^ T cells throughout their response cascade are linked to a tightly regulated metabolic rewiring (Loftus and Finlay, 2016; Pearce and Pearce, 2017). Accordingly, the different stages of T-cell activation and differentiation are characterized by very distinct metabolic profiles, resulting from the diverse needs that T cells must meet in each of those states (Wang and Green, 2012). For example, it has been shown that naïve and memory T cells rely mainly on a catabolic metabolism, including oxidative phosphorylation (OXPHOS) and fatty acid oxidation (FAO), to efficiently support their resting state (Buck et al., 2015; O’Sullivan, 2019). Meanwhile, effector T cells resort to anabolism-promoting pathways, such as aerobic glycolysis (Chang et al., 2013; Ho et al., 2015) or amino acid metabolism (Kelly and Pearce, 2020), to fuel the high energetic and biosynthetic needs imposed by their rapid proliferation and cytokine production machinery. In this sense, disrupting the ability of effector T cells to rewire their metabolism, for example by chronically restricting their access to glucose, can impair their functionality and anti-tumor response (Chang et al., 2015). Meanwhile, supporting their metabolic needs, for example through amino-acid supplementation (Geiger et al., 2016), or promoting their metabolic rewiring to a memory phenotype, by inducing FAO (Pearce et al., 2009), have been shown to increase their tumor-clearing capacity.

In the last years, substantial knowledge on the metabolism of CD8^+^ T cells at different functional endpoint states, namely naïve, effector, or memory cells, has been gained (Kaech and Cui, 2012; Pearce, 2010). However, little is yet known about how the metabolic needs of CD8^+^ T cells dynamically evolve when transitioning between these different phenotypic states (MacIver et al., 2013; Pearce et al., 2013). This dynamic characterization cannot be inferred from time-resolved bulk metabolic measurements because individual T cells will not evolve synchronously throughout their response cascade. Thus, bulk readouts will represent a population average of all different cell states being sampled. Tackling this problem therefore calls for single-cell methodologies, able of capturing and resolving the different metabolic states of all individual cells present in a dynamically-evolving, heterogeneous cell population (Artyomov and Van den Bossche, 2020). A number of recent studies have employed such approaches to characterize dynamic aspects of T-cell metabolism at the single-cell level (Ahl et al., 2020; Hartmann et al., 2021; Levine et al., 2021). Specifically, both (Ahl et al., 2020) and (Hartmann et al., 2021) showed how the rewiring of particular metabolic pathways (such as glycolysis, OXPHOS, or FAO) known to occur during *in vivo* T-cell activation can be recapitulated based on time-resolved mass-cytometry measurements of metabolic enzymes in *in vitro* activated human T cells, while further providing a dynamic description of this rewiring. Using also mass cytometry, but this time based both on *in vitro* and *in vivo* activating and differentiating mouse T cells, (Levine et al., 2021) further leveraged on the power of single-cell measurements to uncover a metabolically distinct and previously unidentified transient state during early T-cell activation, characterized by both high glycolytic and OXPHOS activities. These recent studies highlight the importance of studying the metabolism of T cells from a dynamic viewpoint, focused not only on endpoint states but also on the transitions between them, in order to gain new insights on T-cell metabolism and its link to T-cell responses. However, due to the technical constraints imposed by mass cytometry measurements (Rossi et al., 2021), these studies were still limited to at most a few tens of metabolic enzymes, focusing exclusively on central carbon metabolism. Thus, a comprehensive metabolism-wide dynamic analysis of the metabolic pathways supporting CD8^+^ T cells in their transition between phenotypic states remains missing.

In this work, we used single-cell RNA-sequencing (scRNA-seq) to provide a global description of the dynamic metabolic rewiring of CD8^+^ T cells transitioning *in vitro* through the activation and differentiation cascade. Our analysis recapitulates a broad number of metabolic aspects previously reported for *in vivo* activated and effector/memory differentiated CD8^+^ T cells, highlighting the physiological relevance of our *in vitro* scRNA-seq approach. In addition, we identify multiple dynamic alterations in the metabolism of CD8^+^ T cells not previously described in the literature. One of these transient metabolic dependencies is the synthesis of asparagine via asparagine synthetase (ASNS), whose expression profile increases during T-cell activation, sharply peaks in the effector state, and later drops towards memory formation. We consequently find that transiently interfering with these ASNS expression dynamics during *in vitro* differentiation, by means of asparagine depletion from the environment, leads to a delay in central-memory T-cell polarization. Furthermore, and consistent with our *in vitro* observations, we show that preventing the decay of ASNS by stable overexpression *in vivo* drives effector T-cell expansion and decreases long-term central-memory T-cell formation. In summary, we provide and further validate here a novel, metabolism-wide dynamic description of the metabolic alterations undergone by CD8^+^ T cells during activation and effector/memory differentiation. Aside from its relevance as a resource for hypothesis generation in the context of immunometabolism, we expect our data set to be a useful tool for the benchmarking of scRNA-seq analysis pipelines focused on modeling and studying dynamic processes.

## RESULTS

### Single-cell RNA-seq captures the population dynamics of *in vitro* activating/differentiating CD8^+^ T cells

The T-cell activation and differentiation cascade is a dynamic process, and the different functional states acquired by T cells throughout this cascade are linked to distinct metabolic programs (Wang and Green, 2012). Therefore, T-cell metabolism needs to also change dynamically in order to support T-cell activation and differentiation (MacIver et al., 2013; Pearce et al., 2013). To characterize the dynamics of T-cell metabolism in a defined setting, we adopted a widely established approach for the *in vitro* activation and further polarization of naïve CD8^+^ T cells into effector- or memory-like cells (Cornish et al., 2006; Mueller et al., 2008). Specifically, we relied on a generic *in vitro* stimulation with anti-CD3 (in the presence of anti-CD28 co-stimulation and the pro-proliferative cytokine IL-2), followed by differentiation by addition of either IL-2 (for effector polarization) or the pro-homeostatic cytokine IL-15 (for memory polarization) (Zhang et al., 1998) (**Figure S1A**). Since cellular metabolism is highly dependent on the local nutrient availability (Elia and Fendt, 2016; Rinaldi et al., 2018; Wei et al., 2017), we further used a culture medium resembling the nutrient composition found *in vivo* in human plasma (*blood-like medium*, BLM, **Supplementary Table 1**) (Fernández-García and Fendt, 2019; Tardito et al., 2015), to make our *in vitro* results more representative of an *in vivo* situation. In addition, we isolated naïve CD8^+^ T cells from the spleens of OT-I mice (Clarke et al., 2000; Hogquist et al., 1994), to minimize the impact of polyclonality and ensure that all (metabolic) heterogeneity during the *in vitro* response stemmed from different activation/differentiation states, even under non-antigen-specific stimulation.

scRNA-seq provides a snapshot of the individual gene-expression profiles of all cells in a particular cell population at a given point in time. This static picture can however be used to infer the dynamics of an evolving population, provided that the different states of this evolution are represented in the analyzed sample (Grün, 2018; Stubbington et al., 2017). Thus, we used time-resolved FACS-analysis (based on the surface markers CD69, CD25, CD62L, and CD44) to determine the time points of optimal cell-state heterogeneity during our *in vitro* activation and effector/memory polarization (**Figures S1B-C**). Based on these measurements, we identified the 24h activation (**Figure S1B**) and 144h differentiation (for both effector and memory polarization) time points (**Figure S1C**), as those showing the optimal heterogeneity of cell states. In particular, after 24h of activation we observed at least five distinct populations, evolving from the naïve state (CD69^Lo^ CD25^Lo^ CD62L^Hi^ CD44^Lo^) through transient early and intermediate activation states (characterized by simultaneous upregulation of CD69 and loss of CD62L, followed by CD25 upregulation), until reaching a late activation state (characterized by the eventual upregulation of CD44) (**Figures 1A-C**). Meanwhile, our 144h effector differentiation sample (**Figure 1D**) presented four distinct phenotypes, all of them CD69^Hi^ CD44^Hi^. On the one hand, two larger (Weninger et al., 2001), blasting (**Figure S1D**), IL-2-responsive (CD25^Hi^) populations, which we identified as undifferentiated precursors (*Undiff*, CD62L^Hi^), representing the starting differentiation state (**Figure S1C**), and effector-like cells (*Teff*, CD62L^Lo^) (**Figure 1D**). On the other hand, two smaller (Weninger et al., 2001), non-blasting (**Figure S1D**), IL-2-non-responsive (CD25^Lo^) memory-like populations (Kalia et al., 2010), displaying either a central-memory (CD62L^Hi^) or effector-memory (CD62L^Lo^) phenotype, which we designate as central-memory precursors (*Tcm*_*p*_, CD62L^Hi^) and effector-memory precursors (*Tem*_*p*_, CD62L^Lo^) (**Figure 1D**). Finally, our 144h memory differentiation comprised just two distinct populations, both CD69^Hi^ CD44^Hi^ CD25^Lo^, corresponding to central-memory (*Tcm*, CD62L^Hi^) and effector-memory (*Tem*, CD62L^Lo^) cells (**Figure 1E**).

**Figure 1.**
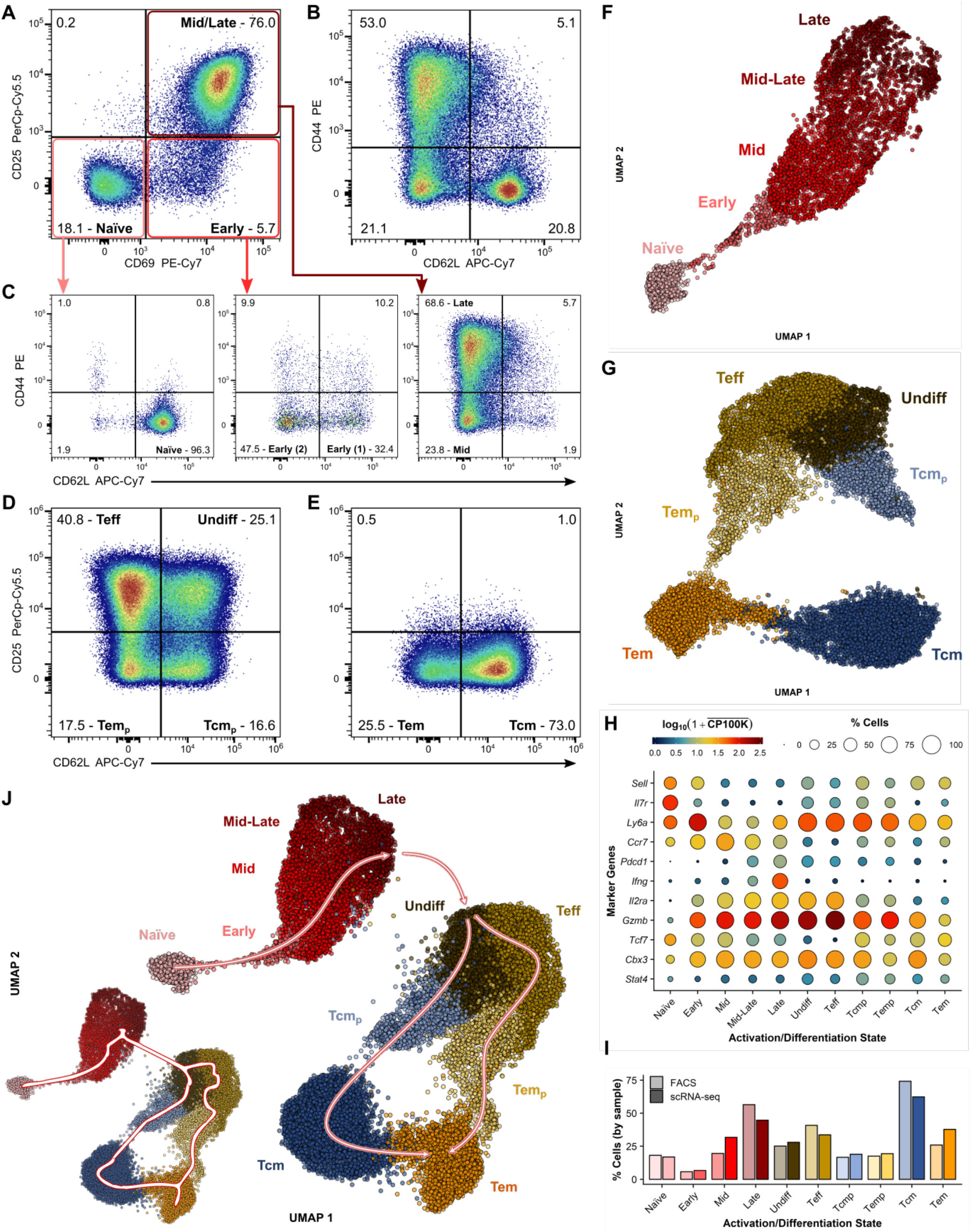
Single-cell RNA-seq captures the population dynamics of *in vitro* activating/differentiating CD8^+^ T cells. **(A–C)** FACS plots of CD25 *vs*. CD69 **(A)** and CD44 *vs*. CD62L **(B)** profiles of stimulated cells at the optimal activation time point (t=24h), identifying at least 5 distinct activation states, based on inspection of the CD44 *vs*. CD62L profiles for the 3 different subpopulations found in the CD25 *vs*. CD69 plane **(C)** **(D–E)** FACS plots of CD25 *vs*. CD62L profiles for effector (IL-2)-polarized (**D**) and memory (IL-15)-polarized (**E**) cells at the optimal differentiation time point (t=144h), identifying at least 4 and 2 distinct states (respectively), based on these surface markers alone. **(F–G)** UMAP plots for the scRNA-seq data corresponding to the 24h activation sample **(F)** and the two 144h differentiation samples combined **(G)**, with assignment of the underlying cell states identified based on the expression profiles of known CD8^+^ T-cell phenotypic markers at the gene expression level. In **(G)**, the upper and lower parts of the UMAP plot correspond respectively to our effector-polarized and memory-polarized samples. **(H)** Gene expression *vs*. cell state profiles for marker genes used to assist cell-state annotation. Log-transformed gene expression levels are indicated by the color scale, where 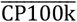 denotes the average gene expression level (in counts per 100k reads) over all cells in a given cell state, while the areas of the circles represent the percentage of cells with non-zero expression of each gene among all cells in each cell state. **(I)** Comparison between the fractions of cells (relative to the total in each sample) assigned to each cell state based on FACS (light bars) or scRNA-seq (dark bars) data. For the FACS data, the *Early* activation state represents the sum of the *Early (1)* and *Early (2)* populations in **(C)**, since these could not be separated based on scRNA-seq. For the scRNA-seq data, the *Late* activation state represents the sum of the *Mid-Late* and *Late* populations in **(F)**, since these could not be separated by FACS analysis. **(J)** UMAP plot for the full merged scRNA-seq data set (all 3 samples combined), with the arrows indicating the expected evolutive trajectory connecting the various cell states based on time-resolved FACS data. The trajectory predicted by Monocle is indicated by the smoothed curve in the inset. The color code for the different cell states matches that used in **(F-G)**. All FACS plots gated on live, single CD8^+^ T cells, corresponding to the merge of 3 replicates, and representative of 4 independent experiments.

We next performed scRNA-seq on cell samples collected at the previously determined optimal time points. Our scRNA-seq data analysis for the three corresponding samples (24h activation, 144h effector polarization, and 144h memory polarization) revealed a continuum of cell states within each of them in the dimensionally-reduced space (UMAP) (**Figures 1F-G**). This is expected from the gradually evolving nature of the activation/differentiation process. To define the underlying cell-state dynamics within this continuum, we resorted to analyzing the expression profiles, in the dimensionally-reduced space, of an array of known CD8^+^ T-cell activation and differentiation marker genes (**Figures S1E-F**), including those encoding the protein surface-markers used in our FACS analysis (**Figure 1H**).

For the 24h activation sample (**Figures 1F, S1E**), we first identified the *Naïve* population (**Figure 1F**, bottom left) based on its higher expression of the naïve markers *Il7r* (CD127) and *Sell* (L-Selectin, CD62L) (Best et al., 2013) (**Figure S1E**). *Sell* expression further extended along the thin streak emerging from the naïve cluster. This, in combination with the high expression detected there for *Ly6a* (**Figure S1E**), a known early activation marker (Best et al., 2013), allowed us to assign this region to *Early* activated cells (**Figure 1F**). *Pdcd1* (PD-1), now widely regarded as a marker of T-cell exhaustion, is also known to be progressively upregulated during T-cell activation (Ahn et al., 2018). The bimodal *Pdcd1* expression profile observed in our data (**Figure S1E**) thus prompted us to subdivide the remaining, larger group of cells on the upper half of the reduced-space projection into *Mid* (*Pdcd1*^Lo^) and late-activation (*Pdcd1*^Hi^) states (**Figure 1F**). Notably, we observed that expression of the chemokine-receptor encoding gene *Ccr7* (CCR7) peaked within the former mid-activation state (**Figure S1E**), being further downregulated towards the late activation one, coinciding, as previously shown in the literature (Bjorkdahl et al., 2003), with the upregulation of genes involved in cytokine production, such as *Ifng* (IFN-γ, **Figure S1E**). In line with this, we could further observe a clear separation between cells expressing either intermediate or high levels of *Ifng* in the late-activation state (**Figure S1E**). Therefore, we further subdivided the latter into a transitional *Mid-Late* activation state (*Ifng*^Int^), and a proper *Late* activation state (*Ifng*^Hi^) (**Figure 1F**), with the latter presenting also a higher expression level of other cytokine-associated genes, such as *Gzmb* (Granzyme B, **Figure 1H**).

In the case of our 144h effector-polarized sample (**Figures 1G, S1F**, top group), *Il2ra* (CD25) expression directly allowed us to tell apart the IL2-responsive (*Il2ra*^Hi^) cells, on the upper half of the UMAP, from the IL2-non-responsive (*Il2ra*^Lo^) memory precursors on the lower half (**Figures 1G, S1F**), with the latter presenting in turn a higher expression of the memory marker *Tcf7* (TCF-1) (Best et al., 2013; Kakaradov et al., 2017) (**Figure S1F**). Based on this first division, *Sell* expression (**Figure S1F**) further allowed us to identify the right branch of the UMAP with undifferentiated (*Undiff, Sell*^Hi^*Il2ra*^Hi^, top-right) and central-memory precursor (*Tcm*_*p*_, *Sell*^Hi^*Il2ra*^Lo^, bottom-right) cells, and to assign their counterparts in the left branch as effector (*Teff, Sell*^Lo^*Il2ra*^Hi^, top-left) and effector-memory precursor (*Tem*_*p*_, *Sell*^Lo^*Il2ra*^Lo^, bottom-left) cells (**Figure 1G**). These assignments were consistent with the expected connectivity between these populations based on our FACS data (**Figure 1D**), and were confirmed by evaluation of other markers, such as *Gzmb*, whose expression level peaked, as expected (Best et al., 2013; Kakaradov et al., 2017), for the effector population (**Figures 1H, S1F**). On the other hand, our 144h memory-polarized sample (**Figures 1G, S1F**, bottom group) displayed a clear division in two large clusters, as expected from our FACS data (**Figure 1E**). *Sell* expression did in this case not allow us to univocally classify these two clusters as either *Tcm* or *Tem* cells, since comparable levels were found in both (**Figure S1F**). However, based on the relative frequency of these cell states expected from our FACS data (**Figure 1E**), and on the proximity of either cluster in the UMAP space to the respective memory-precursor (*Tcmp*/*Temp*) cluster, we could assign the larger cluster on the right-hand side to *Tcm* cells, with the smaller left-hand side cluster thus representing *Tem* cells (**Figure 1G**). We further confirmed this annotation by examining the expression levels of other genes known to be differentially expressed by *Tcm* (e.g. *Cbx3*) or *Tem* (e.g. *Stat4*) cells (Kakaradov et al., 2017; Szabo et al., 2019) (**Figures 1H, S1F**).

We next proceeded to validate our cell state assignments using the corresponding FACS data and additional scRNA-seq analysis pipelines. We found a strong agreement between the sample-specific fractions of each of the 11 distinct cell states inferred from the individual scRNA-seq data sets (5 for 24h activation: *Naïve, Early, Mid, Mid-Late*, and *Late*; 4 for 144h effector polarization: *Undiff, Teff, Tcm*_*p*_, and *Tem*_*p*_; and 2 for 144h memory polarization: *Tcm* and *Tem*), and those derived from FACS measurements of the corresponding samples (**Figure 1I**). Our cell state assignment was additionally confirmed by applying Monocle-based trajectory inference (Cao et al., 2019), in the dimensionally-reduced space, to the combination of the three data sets, which accurately captured the connectivity between the different cell states expected based on FACS analysis (**Figure 1J**). Furthermore, we confirmed the expected directionality of the transitions between the different states in each sample based on RNA velocity (Bergen et al., 2020; La Manno et al., 2018) calculations (**Figure S1G**). In summary, we demonstrate that the population dynamics of *in vitro* activating and differentiating cells can be accurately captured based on single-time point scRNA-seq measurements.

### Single-cell RNA-seq identifies known aspects of *in vivo* CD8^+^ T-cell metabolism during activation and differentiation

To date, changes in CD8^+^ T-cell metabolism have been best characterized during the transition from a naïve to an activated state (Pearce and Pearce, 2013). We thus first used our 24h activation sample to (a) validate the fidelity of our *in vitro* model in recapitulating known aspects of *in vivo* T-cell metabolism during activation, and (b) benchmark the potential of our scRNA-seq data analysis pipeline to capture the underlying activation dynamics in these metabolic pathways. For this, we used gene-set variation analysis (GSVA) (Hänzelmann et al., 2013) to determine per-cell gene-signature scores for several pathways whose activity has been well characterized in the past in both the naïve and fully activated cell states. We find a strong agreement between the results of our pathway analysis (**Figure 2A**) and several known key aspects of CD8^+^ T-cell metabolism and signaling upon activation. For example, we confirm the previously described PI3K/Akt/mTOR/Myc-orchestrated (Chi, 2012; MacIver et al., 2013; Waickman and Powell, 2012) switch from a markedly oxidative metabolism (driven by OXPHOS and FAO) (Wang et al., 2011) to a biosynthetic program driven by aerobic glycolysis (Cham and Gajewski, 2005; Cham et al., 2008), glutamine uptake and catabolism (Carr et al., 2010; Wang et al., 2011), and fatty-acid synthesis (FAS) (Kidani et al., 2013), accompanied by an upregulation of the cells’ cytokine-production machinery upon activation (**Figure 2A**). In addition, we also confirm more recent observations in less commonly studied pro-anabolic pathways shown to be upregulated by T cells in response to TCR stimulation, such as polyamine biosynthesis (Wang et al., 2011), the methionine cycle (Sinclair et al., 2019), and the mevalonate pathway for cholesterol synthesis (Thurnher and Gruenbacher, 2015) (**Figure 2A**). Importantly, aside from capturing the known changes in these pathways between the naïve and activated endpoint states, our data further provide a dynamic description of these changes (**Figure 2B**). In line with this, our results recapitulate recent observations from *in vivo* ^13^C-labeling (Ma et al., 2019) and *in vivo* single-cell mass-cytometry data (Levine et al., 2021) demonstrating that, contrary to the widespread notion, both glycolysis and OXPHOS are important in activated CD8^+^ T cells *in vivo*, with OXPHOS also being rapidly upregulated during early activation (Levine et al., 2021) (**Figure 2B**), and glycolysis taking over only at later activation stages (but not at the expense of OXPHOS activity) (**Figure 2B**). In summary, our *in vitro* pathway analysis strategy is able to identify known aspects of the *in vivo* metabolic rewiring undergone by CD8^+^ T cells during activation, while further providing a novel dynamic description of this rewiring.

**Figure 2.**
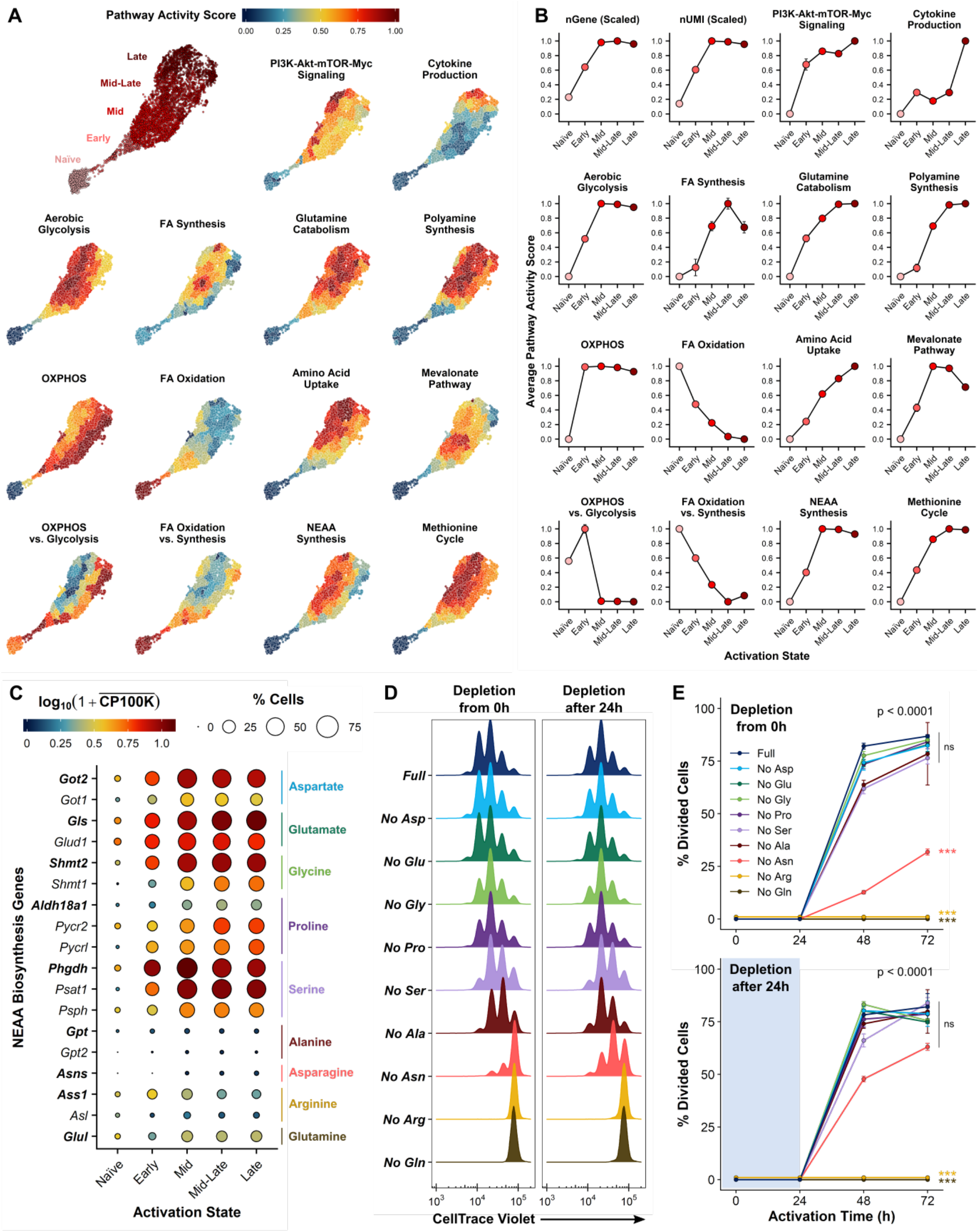
Single-cell RNA-seq identifies known aspects of *in vivo* CD8^+^ T-cell metabolism during activation and differentiation and predicts the dependence of activating CD8^+^ T cells on amino acid uptake *vs*. synthesis. **(A)**UMAP plots for the scRNA-seq data corresponding to the 24h activation sample, color-coded based on GSVA-based pathway activity scores for metabolic/signaling pathways known to be up/down-regulated along CD8^+^ T-cell activation. Pathway activity scores are averaged over all cells in a given fine-grained cluster, and further scaled to the range 0–1 for each individual pathway (see Methods). A replica of Figure 1F is included for reference. **(B)**Cell state-averaged dynamic pathway-activity profiles during CD8^+^ T-cell activation, corresponding to the metabolic and signaling pathways in **(A)**. Pathway activity scores are averaged over all cells in each activation state, and further scaled to the range 0–1 for each individual pathway. Cell-state averaged dynamic profiles for the number of genes (nGene) and library size (nUMI), scaled relative to their maximum value, are included in the first two panels for reference. The various activation states are indicated both in the *x* axis and via the dot color code. Error bars represent standard errors of the mean (±SEM). **(C)**Gene expression *vs*. cell state profiles for key genes involved in non-essential amino acid (NEAA) biosynthesis. Log-transformed gene expression levels are indicated by the color scale, where 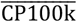 denotes the average gene expression level (in counts per 100k reads) over all cells in a given cell state, while the areas of the circles represent the percentage of cells with non-zero expression of each gene among all cells in each cell state. Genes encoding the rate-limiting enzymes for the synthesis of the various NEAAs are indicated in bold face, with the corresponding NEAAs indicated on the right-hand side. **(D)**CellTrace^®^ Violet dilution profiles after 48h of *in vitro* stimulation in nutrient-replete BLM (Full), or upon single-NEAA depletion, either from the start of the stimulation **(left)**, or after 24h stimulation in nutrient-replete BLM **(right)**. Data correspond to averages of 4 independent culture wells per condition, and are representative of 2 independent experiments. **(E)**FACS-based time-course profiles for the percentage of divided cells (determined from CellTrace^®^ Violet dilution) over a total of 72h of stimulation, for identical NEAA depletion conditions as in **(D)**. Data correspond to averages of 4 independent culture wells per condition and time point, and are representative of 2 independent experiments. Error bars represent standard deviations (±SD). Statistical analyses based on F-Tests on 2^nd^ degree polynomials (see Methods), with global F-Test p-values shown on the top-right corners, and pairwise F-Test significance levels relative to the Full condition displayed next to the profiles for the different conditions.

We next decided to extend our pathway analysis approach to our 144h effector and memory differentiation samples (**Figure S2A**). As expected from our current knowledge of the *in vivo* metabolism of effector *vs*. memory CD8^+^ T cells (van der Windt and Pearce, 2012), we observed a downregulation of the biosynthetic program acquired during activation along the transition from an undifferentiated or effector state to a memory phenotype (**Figure S2A**). This switch is best evidenced by the increase in the relative importance of catabolic *vs*. anabolic pathways, namely OXPHOS *vs*. glycolysis or FAO *vs*. FAS, upon memory differentiation, and is further accompanied, as expected (Cui and Kaech, 2010), by a reduction in the cytokine-producing activities of the cells (**Figure S2A**). Our data further captured the known higher reliance on OXPHOS of *Tcm vs. Tem* cells (van der Windt et al., 2012), and supported recent observations showing that long-lived *Tcm* cells rely on intrinsic FAS to fuel their FAO (Cui et al., 2015; O’Sullivan et al., 2014) (**Figure S2A**). Notably, our data also suggest that, much like during activation both glycolysis and OXPHOS are upregulated, with glycolysis taking over only upon full activation (Levine et al., 2021), the downregulation of glycolysis in the transition to a memory phenotype seems to be concomitant with a downregulation in OXPHOS, with OXPHOS only becoming dominant for fully differentiated (*Tcm/Tem*) memory cells (**Figure S2A**). In this sense, the sustained glycolytic activity observed in the memory precursors of our effector-polarized sample (*Tcm*_*p*_*/Tem*_*p*_) (**Figure S2A**) may partly be due to the presence of IL-2 stimulation, in agreement with the known role of the latter in promoting glycolysis (Bauer et al., 2004). Interestingly, our data also suggest that, aside from OXPHOS and fatty-acid metabolism (including both FAO and FAS), glutamine catabolism may also be an important source of carbon to drive the oxidative program of *Tcm* cells, and that *Tcm* and *Tem* cells may differentially rely on methionine metabolism (**Figure S2A**). In conclusion, just like in the case of activation, our *in vitro* pathway analysis approach is able to identify the known changes in *in vivo* CD8^+^ T-cell metabolism during effector/memory differentiation, while also providing new insights on the differential activity of previously unexplored pathways in different memory T-cell subsets.

Finally, it is important to note that a key factor enabling our *in vitro* data to recapitulate metabolic aspects representative of an *in vivo* situation is the use of a physiological culture medium. This is clearly illustrated, for example, in the case of pyruvate utilization, for which notable differences between *in vivo*- and *in vitro*-activated CD8^+^ T cells were recently reported (Ma et al., 2019). Specifically, it was shown that, whereas *in vitro*-activated CD8^+^ T cells preferentially shunt pyruvate into the TCA cycle via pyruvate carboxylase, *in vivo* activated CD8^+^ T cells rather oxidize it to generate mitochondrial acetyl-CoA using pyruvate dehydrogenase (Ma et al., 2019). In line with the *in vivo* rather than *in vitro* data in (Ma et al., 2019), we observe high expression levels for genes encoding proteins in the pyruvate dehydrogenase complex (e.g. *Pdha1, Pdhb*), coupled to a marked absence of *Pcx* (pyruvate carboxylase) expression, in a majority of our *in vitro*-activated and differentiated CD8^+^ T cells, regardless of their activation or differentiation state (**Figure S2B**). This discrepancy with prior *in vitro* data (Ma et al., 2019) is likely due to the fact that both glucose and pyruvate are at physiological levels in our culture medium, while being 2-5 times (glucose) and 10 times (pyruvate) higher in usual T-cell culture media (Ma et al., 2019) (**Supplementary Table 1**). This highlights once more the importance of using media resembling physiological conditions for a more faithful representation of cellular metabolism in *in vitro* studies (Cantor et al., 2017; Leney-Greene et al., 2020; Vande Voorde et al., 2019).

### Single-cell RNA-seq predicts the dependence of activating CD8^+^ T cells on amino acid uptake *versus* synthesis

The importance of amino acid metabolism during CD8^+^ T-cell responses is now widely acknowledged (Kelly and Pearce, 2020). Recent studies have shown that CD8^+^ T-cell activation relies not only on the uptake of essential amino acids (Ananieva et al., 2016; Sinclair et al., 2019), but also on the extracellular supply of various non-essential amino acids (NEAAs) such as glutamine (Carr et al., 2010; Wang et al., 2011), arginine (Choi et al., 2009), serine (Ma et al., 2017), alanine (Ron-Harel et al., 2019), or asparagine (Hope et al., 2020; Torres et al., 2016; Wu et al., 2021). Thus, despite their non-essential nature, the biosynthetic machinery of activating CD8^+^ T cells alone cannot meet the requirements for some NEAAs, rendering them as virtually essential from the point of view of T-cell activation. Interestingly, using our pathway analysis strategy, we also identified both amino-acid biosynthesis and uptake as hallmarks of CD8^+^ T-cell activation (**Figure 2A**). We thus asked whether the dependence of activating CD8^+^ T cells on the uptake *vs*. synthesis of different nutrients may be predicted from the dynamic expression profiles of the genes regulating their production. To answer this question, we first compared the dynamic expression profiles, based on our 24h activation scRNA-seq data, for individual genes involved in NEAA biosynthesis (**Figure 2C**). From this comparison, we detected a clear separation in two groups. On the one hand, we found the genes controlling the synthesis of aspartate, glutamate, glycine, proline, and serine to be rapidly upregulated during T-cell activation (**Figure 2C**, top). This suggests that, provided the corresponding substrates, activating CD8^+^ T cells may be able to cope with their requirements for these amino acids based on synthesis alone. On the other hand, we found a sustained low expression during activation for the genes responsible for alanine, asparagine, arginine, and glutamine synthesis (**Figure 2C**, bottom). This, in turn, suggests that CD8^+^ T-cell activation may rely predominantly on the uptake of these amino acids from the extracellular environment, as reported in the literature.

To validate the predictions from our scRNA-seq data, we designed an *in vitro* NEAA depletion screen in which we independently removed either of the above 9 amino acids from culture from the beginning of activation (**Figure S2C**). Consistent with our predictions, and in agreement with prior studies, we found that depletion of either glutamine, arginine, or asparagine greatly impaired activation-induced proliferation (**Figures 2D-E, S2D**) and cell growth (**Figure S2E**), as well as surface expression of the late-activation marker CD44 (**Figure S2F**). This impairment was particularly significant in the case of glutamine and arginine, in the absence of which proliferation was totally ablated, and was further evident based on cell morphology, with cells lacking these 3 amino acids closely resembling unstimulated (naïve) cells after 48h of activation (**Figure S2G**). Conversely, and again in line with our predictions, depletion of aspartate, glutamate, glycine, or proline had no significant impact in CD8^+^ T-cell activation (**Figures 2D-E, S2D-F**), whereas serine depletion led only to a minor reduction in proliferation (**Figures 2D-E, S2D**). The latter is consistent with the *in vitro* observations of (Ma et al., 2017), where extracellular serine depletion resulted predominantly in a decrease in CD4^+^ rather than CD8^+^ T-cell proliferation, and only when cells were simultaneously deprived from glycine. It is further in agreement with more recent observations showing how both serine depletion and serine biosynthesis inactivation are required to significantly ablate CD8^+^ T-cell expansion *in vivo* (Ma et al., 2019). Finally, and aside from a minor proliferation impairment comparable to that caused by serine deprivation (**Figures 2D-E, S2D**), alanine depletion led only to a slight delay in cell growth and CD44 expression (**Figures S2E-F**), from which cells recovered after 48h of activation. This suggests that CD8^+^ T cells, unlike their CD4^+^ counterparts investigated in (Ron-Harel et al., 2019), may be able to upregulate alanine transaminase (GPT) expression in response to alanine deprivation, highlighting the importance of considering that metabolic differences exist between T-cell subtypes (Jones et al., 2017; Ma et al., 2017).

Next, we asked how these dependencies change during activation. Thus, we implemented an analogous NEAA depletion screen, but removing the corresponding amino acids only after 24h of activation (**Figure S2C**). Consistent with our observations upon depletion from the beginning of stimulation, depletion of either aspartate, glutamate, glycine, proline, serine, or alanine after 24h of activation had overall no significant impact in CD8^+^ T-cell activation (**Figures 2D-E, S2D-F**). Arginine and glutamine depletion after 24h of activation, on the other hand, still led to a noticeable activation impairment (**Figure S2G**), with cells being unable to proliferate at all in the absence of either amino acid (**Figures 2D-E, S2D**), and both cell growth and CD44 expression being severely hindered under glutamine deprivation (**Figures S2E-F**). Intriguingly, asparagine depletion after 24h of activation had only a minor impact in cell proliferation (**Figures 2D-E, S2D**) and growth (**Figure S2E**), but otherwise no significant impact on CD44 expression (**Figure S2F**), with the resulting cell morphology resembling that of asparagine-replete conditions (**Figure S2G**). Our observations are in agreement with recent reports suggesting that CD8^+^ T cells may be able to upregulate asparagine synthetase (ASNS) expression in response to asparagine depletion during the later stages of activation (thus enabling them to function independently of the extracellular supply from then on), but not during the very early activation stages (thus failing to exit the naïve state under complete absence of extracellular asparagine) (Hope et al., 2021). In summary, we show that both NEAA uptake and synthesis are hallmarks of CD8^+^ T-cell activation, and that the differential dependence of these cells on uptake *vs*. synthesis for different NEAAs can be predicted based on single-cell gene expression measurements.

### Single cell RNA-seq identifies the dynamic orchestration of CD8^+^ T-cell metabolic programs during activation and differentiation

In order to further benchmark the potential of our scRNA-seq-based approach for capturing dynamic aspects of CD8^+^ T-cell metabolism on a broad scale, we decided to focus our attention on polyamine metabolism. Polyamine metabolism is implicated in a variety of cellular processes driving cell growth, proliferation, and homeostasis, such as DNA replication, apoptosis, and autophagy (Pegg, 2016). As such, it has received considerable attention over the past years in the context of cancer (Arruabarrena-Aristorena et al., 2018). Nevertheless, polyamine metabolism is still relatively unexplored in the context of CD8^+^ T cells (Puleston et al., 2017), where our current knowledge is mainly limited to the importance of Myc-driven polyamine biosynthesis during T-cell activation (Wang et al., 2011). The study of polyamine metabolism is, moreover, particularly interesting from a systems level perspective. This is because it involves the activities of various upstream (e.g. Myc, but also mTOR) and downstream (e.g. eIF5A, via its hypusination) signaling pathways, as well as of multiple ancillary metabolic pathways, including polyamine biosynthesis itself, but also glutamine, proline, and arginine metabolism, or the methionine and urea cycles (Arruabarrena-Aristorena et al., 2018; Puleston et al., 2017). We thus asked whether our scRNA-seq data would be able to identify the dynamics underlying the orchestration of these multiple pathways, and whether this would allow us to gain new insights on the role polyamine metabolism in yet unexplored contexts such as CD8^+^ T-cell differentiation.

To answer these questions, we started by examining the dynamic expression profiles of some of the key genes involved in polyamine metabolism (Puleston et al., 2017). Specifically, we focused on the main four genes driving polyamine synthesis, namely *Odc1* (ornithine decarboxylase), *Srm* (spermidine synthase), *Sms* (spermine synthase), and *Amd1* (s-adenosylmethionine decarboxylase), the latter required for the production of decarboxylated s-adenosyl methionine (SAM), a necessary cofactor in spermidine and spermine synthesis. Strikingly, we found that the dynamic expression profiles of these four genes not only displayed similar trends, but seemed moreover tightly coordinated with each other throughout both activation and differentiation (**Figure 3A**, top half). Conversely, the dynamic profiles of genes encoding enzymes opposing polyamine synthesis, such as *Paox*/*Smox* (polyamine/spermine oxidase, responsible for polyamine catabolism), or *Ass1*/*Asl* (argininosuccinate synthase/lyase, driving the competing urea cycle), were clearly inversely-correlated with the former along the entire activation and differentiation cascade (**Figure 3A**, bottom half). We quantitatively confirmed this based on the correlation coefficients between the expression levels of these genes and those of *Odc1*, the rate-limiting step in the synthesis pathway, considering all different cell states identified by our analysis (**Figure S3A**). Overall, this suggests the existence of a strong dynamic coordination between the activities of the multiple pathways tied to polyamine metabolism throughout the entire CD8^+^ T-cell response cascade.

**Figure 3.**
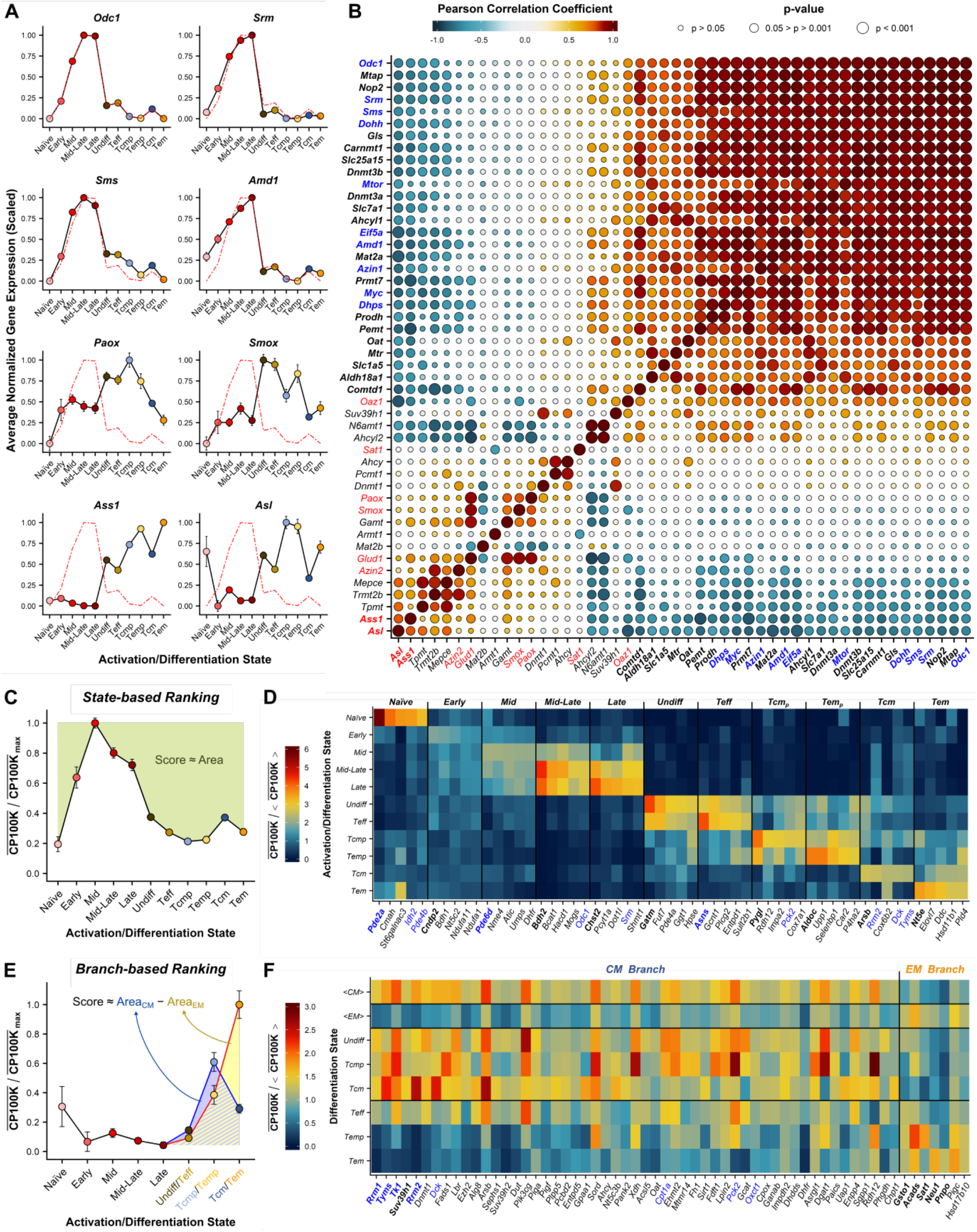
Single-cell RNA-seq captures the dynamic orchestration of CD8^+^ T-cell metabolic programs during activation and differentiation and identifies novel dynamic metabolic dependencies. **(A)**Cell state-averaged expression profiles for select genes involved in polyamine synthesis **(top half)** or in polyamine catabolism and the urea cycle **(bottom half)**, with the red-dashed line showing the profile for *Odc1*, for comparison purposes. Gene expression levels are first averaged over all cells in each state, and the resulting expression *vs*. cell-state profiles are scaled to the range 0–1 for each gene, by subtracting the average expression level for the lowest expressing state, and then dividing by the difference in average expression between the highest and lowest expressing states (see Methods). The various activation/differentiation states are indicated both in the *x* axis and via the dot color code. Error bars represent standard errors of the mean (±SEM), considering the expression levels of all cells in each state. **(B)**Heatmap plot with mutual correlation coefficients between the expression profiles of a variety of genes related to polyamine metabolism. The color scale represents the Pearson correlation coefficient (*R*) between every pair of genes, while the dot sizes represent the corresponding p-values. Genes are sorted by their *R* values relative to *Odc1*, with genes with |*R*| *> 0*.*7* indicated in bold face. Genes involved in polyamine synthesis and hypusination, or being known upstream regulators or downstream targets of the latter, are highlighted in blue, while genes opposing polyamine synthesis are highlighted in red. **(C)**Schematic depiction of our *state-based* ranking approach. The magnitude of the green area over the curve, based on the cell-state averaged expression profile normalized relative to the maximum expression level, represents the scoring metric. **(D)**Heatmap plot of cell state-averaged expression profiles for the 5 top-ranked genes for each cell state according to our *state-based* ranking. Scaled expression levels are indicated by the color scale, where 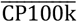 denotes the average gene expression level (in counts per 100k reads) over all cells in a given cell state, and 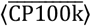 the average of the latter over all cell states. The top-ranked gene for each state is indicated in bold face, with genes discussed in the text highlighted in blue. **(E)**Schematic depiction of our *branch-based* ranking approach. The magnitude of the difference between the blue and yellow areas, based on the cell-state averaged expression profile normalized relative to the maximum expression level, represents the scoring metric. Positive area differences denote genes dominating in the *Tcm*-generating branch, while negative area differences denote genes dominating in the *Tem*-generating branch. **(F)**Heatmap plot of cell state-averaged expression profiles for the 60 top-ranked genes (overall) according to our *branch-based* ranking. Scaled expression levels are indicated by the color scale, where 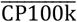 denotes the average gene expression level (in counts per 100k reads) over all cells in a given cell state, and 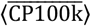 the average of the latter over all cell states (including the activation states, not shown in the heatmap plot). The y-axis entries *<CM>* and *<EM>* represent averaged values over the three states constituting either of the *Tcm*-generating or *Tem*-generating branches, respectively, with their difference being indicative of the scoring metric. The 5 top-ranked genes for each branch are indicated in bold face, with genes discussed in the text highlighted in blue.

To further explore this hypothesis, we subsequently extended our analysis to a larger set of genes tied to polyamine metabolism. This set included not only the former biosynthetic or catabolic genes, but also the known transcriptional regulators/targets of the synthesis pathway, as well as genes participating in ancillary metabolic pathways and hypusine synthesis (Arruabarrena-Aristorena et al., 2018; Puleston et al., 2017). For this, we correlated the dynamic expression profiles of each of those genes against that of *Odc1* (**Figure S3B**), and also against those of all other genes in the set (**Figure 3B**). In agreement with our hypothesis, we observed a high degree of mutual correlation between the dynamic expression profiles of a majority of genes known to be positive regulators or targets of polyamine synthesis. This included, among others, the pathway drivers *Myc* and *Mtor*, as well as the downstream hypusination target *Eif5a*, and the genes responsible for hypusine production *Dhps* and *Dohh* (**Figure 3B**). It further included genes involved in the methionine cycle, such as *Ahcyl1*, the adenosyl/methyl transferases *Mtr, Mat2a*, or *Dnmt3a/b*, necessary for SAM production (prior to its decarboxylation via *Amd1*), and *Mtap*, responsible for methionine salvage from the by-products of polyamine synthesis (**Figure 3B**). Finally, it also included genes involved in ornithine uptake (*Slc7a1*), or its production from extracellular glutamine and/or proline (*Slc1a5, Aldh18a1, Prodh, Oat*) and its further shunting to the cytosol (*Slc25a15*) for polyamine synthesis (**Figure 3B**). Meanwhile, we also found a high degree of mutual correlation, and a simultaneous negative correlation relative to the former genes, for genes known to negatively regulate the synthesis pathway. This included not only the previously mentioned polyamine catabolism and urea cycle genes, but also *Glud1* (glutamate dehydrogenase), diverting glutamate to α-ketoglutarate, rather than to pyrroline-5-carboxylate as required for ornithine synthesis (**Figure 3B**). Interestingly, we could not detect expression for either arginase isoform *Arg1*/*2* or for the agmatinase *Agmat*, while *Azin2* (arginine decarboxylase) expression was again negatively correlated with that of the polyamine biosynthetic genes (**Figure 3B**). This supports prior reports that arginine is not metabolized into ornithine or putrescine during CD8^+^ T-cell activation, with activating cells shunting instead glutamine/proline-derived ornithine to polyamine production (Wang et al., 2011), also in agreement with our results (**Figure 3B**). Thus, our data confirm the existence of a dynamically regulated, joint orchestration of the activities of the various pathways converging in polyamine metabolism in CD8^+^ T cells. More importantly, we show that this orchestration is not limited to CD8^+^ T-cell activation (Wang et al., 2011), but rather extends to the entire CD8^+^ T-cell activation and differentiation cascade. In this respect, our data indicate the existence of a previously unexplored rewiring in CD8^+^ T-cell metabolism, whereby CD8^+^ T cells undergo a switch from polyamine synthesis during activation to polyamine catabolism during differentiation, with this switch being accompanied by an upregulation in their urea cycle activity (**Figures 3A-B, S3B**). This rewiring has a number of potential implications, particularly with regards to arginine metabolism. Indeed, as previously reported in the literature (Choi et al., 2009) and further confirmed by our data (**Figure 2**), CD8^+^ T-cell activation relies heavily on arginine uptake rather than synthesis. Thus, one could speculate that the lower demand for polyamine synthesis during differentiation may allow CD8^+^ T cells to upregulate the activity of the competing urea cycle for arginine synthesis, without further conversion to ornithine. This would in turn enable differentiating CD8^+^ T cells to function independently of an extracellular arginine supply, or alternatively promote their maintenance of elevated intracellular arginine levels, which have been shown to increase T-cell survival and memory formation (Geiger et al., 2016). In summary, we demonstrate how our scRNA-seq-based approach is able to capture the dynamic orchestration of the activities of multiple signaling and metabolic pathways driving CD8^+^ T-cell progression through the activation and differentiation cascade. Moreover, we show how this dynamic information can provide new insights and drive hypothesis generation on previously unexplored aspects of CD8^+^ T-cell metabolism.

### Single cell RNA-seq identifies novel dynamic metabolic dependencies of CD8^+^ T cells

Having validated the ability of our scRNA-seq approach to provide new dynamic insights on known aspects of CD8^+^ T-cell metabolism, we decided to go one step further, and exploit the wealth of information provided by our data to uncover yet unknown dynamic metabolic dependencies of CD8^+^ T cells. To do so, we adopted an untargeted, gene-centric approach (i.e. without a specific focus on pre-defined pathways), taking advantage of the dynamic cell-state resolution provided by our single-cell data. Specifically, we aimed at identifying metabolic genes with expression levels peaking at specific intermediate states along the CD8^+^ T-cell response cascade. The reason is that, due to their transient characteristics, the importance of these genes (and thus of their associated pathways) in particular stages of the CD8^+^ T-cell response may have been previously overlooked based on bulk, endpoint measurements. Additionally, we aimed at also identifying genes whose expression may dictate the fate of CD8^+^ T cells with respect to acquiring a central-memory *vs*. an effector-memory phenotype. This distinction is important, given that the former is a more desirable outcome from a long-term protection, therapeutic point of view (Liu et al., 2020).

For this untargeted analysis, we first restricted our data set to consider only metabolic genes included in the KEGG metabolic pathway database (Kanehisa and Goto, 2000). We further filtered this set of metabolic genes to include only those with expression levels above a predefined threshold in at least one of the 11 activation and differentiation cell states identified in our scRNA-seq data (**Figures 1F-G**), resulting in a total of 750 metabolic genes. Based on this final set of genes, we then implemented two independent ranking metrics. On the one hand, we ranked these genes based on the magnitude of their expression in either of the above mentioned 11 activation and differentiation cell states relative to all other states (**Figure 3C**). This *state-based* ranking (**Figure 3D**) thus tells us to which extent the dynamic upregulation of a particular metabolic gene may be critical in a specific stage of CD8^+^ T-cell activation or differentiation. On the other hand, we ranked these genes based on their differential expression along the *Tcm*-generating “branch” of our *in vitro* differentiation (comprising the *Undiff, Tcm*_*p*_, and *Tcm* states) relative to that along its *Tem*-generating counterpart (comprising the *Teff, Tem*_*p*_, and *Tem* states) (**Figure 3E**). This *branch-based* ranking (**Figure 3F**) thus tells us whether the expression of a particular metabolic gene may dictate the potential of CD8^+^ T cells differentiating into either central-memory or effector-memory cells.

Among the top hits identified in our *state-based* (**Figure 3D**) or *branch-based* (**Figure 3F**) rankings, we found, as expected, several genes whose role in particular stages of CD8^+^ T-cell responses has been described in the past. These include, for example, *Idh2* (mitochondrial isocitrate dehydrogenase), responsible for the rate-limiting step of the TCA cycle, and which we find as the 4^th^ top-ranked gene for the *Naïve* state (**Figure 3D**), in which T cells are known to chiefly rely on oxidative metabolism (Wang et al., 2011). Moreover, we find the polyamine biosynthetic genes *Odc1* and *Srm* ranked 5^th^ and 4^th^, respectively, in the *Mid-Late* and *Late* activation states (**Figure 3D**), in which polyamine synthesis is known to become increasingly important (Wang et al., 2011). We also find *Cpt1a* (carnitine palmitoyl transferase 1α), responsible for the rate-limiting step in β-oxidation, among the top 30 genes for the *Tcm*-generating branch (**Figure 3F**), consistent with the importance of FAO for long-term memory development (van der Windt et al., 2012). Finally, we find *Pck2*, encoding the mitochondrial isoform of *PEPCK* (phosphoenolpyruvate carboxykinase) among the top-ranked genes for both the *Tcm*_*p*_ central-memory precursor state (**Figure 3D**) and the *Tcm*-generating branch (**Figure 3F**), in which we also find *Oxct1* (3-oxoacid CoA-transferase 1), responsible for the rate-limiting step of ketolysis (**Figure 3F**). In line with this, a role for ketone-body driven PEPCK activation in long-term memory formation was recently uncovered in the literature (Ma et al., 2018; Zhang et al., 2020). Thus, it is tempting to speculate that *Oxct1* may serve as a buffer, by diverting the excess ketone-body (particularly, β-hydroxybutyrate) accumulation in *Tcm* cells towards acetyl-CoA production to feed the TCA cycle, in agreement with the observed increase in β-hydroxybutyrate carbon contribution to TCA cycle intermediates in memory T cells shown by (Zhang et al., 2020).

These are just a few examples serving to illustrate the power of our untargeted approach in identifying key metabolic dependencies of CD8^+^ T cells along the activation and differentiation cascade. Importantly, numerous other metabolic genes topping our *state-based* (**Figures 3D, S3C**) and *branch-based* (**Figures 3F, S3D**) rankings, and the metabolic pathways in which they are involved, are still largely unexplored in the context of CD8^+^ T cells, opening up an array of possibilities for hypothesis generation. Among them we found, for example, a variety of cGMP-dependent phosphodiesterase-coding genes (such as *Pde2a, Pde4b*, or *Pde6d*) among the top-ranked genes for the *Naïve* and *Early/Mid* activation states (**Figure 3D**). Some of these phosphodiesterases have been previously linked to the regulation of mitochondrial morphology and ATP production (Monterisi et al., 2017), suggesting a potential relation to the elevated OXPHOS activity observed in early-activated CD8^+^ T cells (Levine et al., 2021). Additionally, we found several genes involved in purine and pyrimidine metabolism (including *Rrm1/2/2b, Tyms, Tk1*, or *Dck*, among others) to be highly ranked in the *Tcm*-generating branch relative to its *Tem*-generating counterpart (**Figure 3F**), with a number of these being also among the top-ranked genes for the *Tcm* state (**Figure 3D**). Thus, one may speculate as to whether nucleotide metabolism could be a previously unexplored hallmark of long-term central-memory formation, consistent with the reported higher self-renewal capacity of *Tcm* cells relative to *Tem* cells (Marzo et al., 2005; Wherry et al., 2003). This, in turn would explain the skewing of our *branch-based* ranking towards the *Tcm*-generating branch (**Figure S3D**), based on the direct relation existing between proliferative capacity and transcriptional activity (Marguerat et al., 2012). Finally, we were intrigued to find *Asns* (asparagine synthetase) as the top-ranked gene for the effector (*Teff*) state (**Figure 3D**), given the markedly low expression levels we had observed for the latter during CD8^+^ T-cell activation (**Figure 2C**). Interestingly, while the role of asparagine metabolism in T-cell activation has been recently reported (Hope et al., 2020; Torres et al., 2016; Wu et al., 2021), its potential impact on CD8^+^ T-cell differentiation, and specifically on memory formation, remains unexplored (Raynor and Chi, 2021). Therefore, we decided to further focus our attention on it.

### The expression dynamics of asparagine synthetase dictate the outcome of T-cell differentiation

*Asns* encodes the enzyme asparagine synthetase (ASNS), responsible for the synthesis of asparagine from aspartate in an ATP-dependent transamidation reaction involving also the conversion of glutamine to glutamate (**Figure S4A**). Based on our *state-based* ranking analysis described above, we identified *Asns* as the top-ranked gene for the effector (*Teff*) state of CD8^+^ T-cell differentiation (**Figure 3D**). We thus first wanted to assess whether our *in vitro* scRNA-seq-based *Asns* expression profile (**Figure S4B**) is representative of that observed during an *in vivo* immune response. Therefore, we isolated naïve CD8^+^ T cells from P14 transgenic mice, in which CD8^+^ T cells express a T-cell receptor recognizing gp33 peptide (an epitope of lymphocytic choriomeningitis virus) loaded MHCI, and then adoptively transferred them into wild type C57BL/6J mice followed by infection with lymphocytic choriomeningitis virus (LCMV) Armstrong strain (**Figure S4C**). We then FACS-sorted CD8^+^ T cells from the spleens of the recipient mice at different time points during the ensuing primary immune response, and measured their relative *Asns* mRNA levels (**Figure S4C**). Strikingly, we found a remarkable agreement between our *in vitro* (**Figure S4B**) and *in vivo* (**Figure 4A**) data, with the latter displaying again a marked upregulation in *Asns* expression during the early effector response, followed by a decay in the transition towards a memory state (**Figure 4A**). Our data thus indicate that, both *in vivo* and *in vitro*, effector CD8^+^ T cells, unlike their memory counterparts, rely heavily on asparagine synthesis. This in turn suggests that, aside from its known impact during early CD8^+^ T-cell activation, ASNS expression levels may play a role in modulating the progression of CD8^+^ T cells through the differentiation cascade.

**Figure 4.**
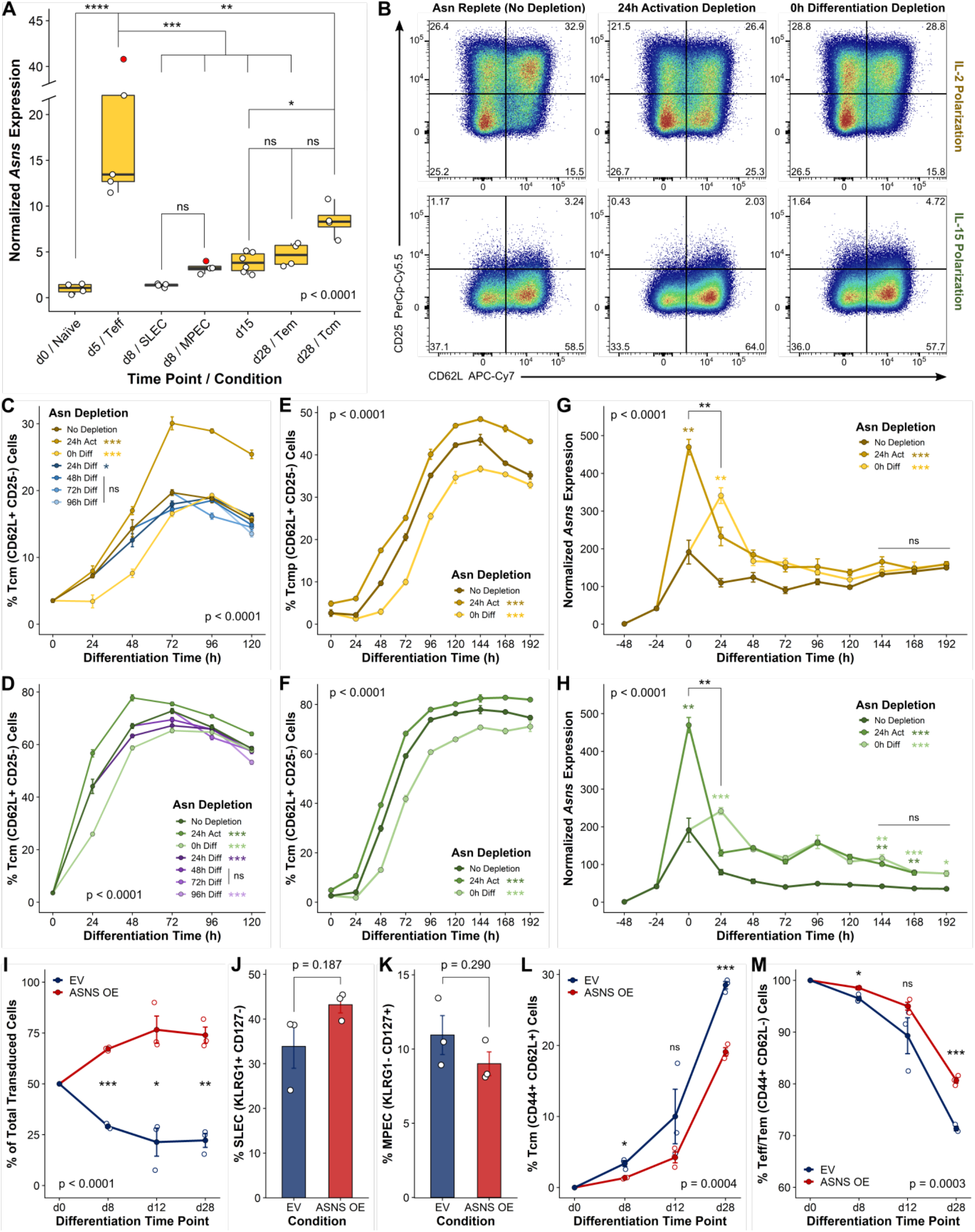
The expression dynamics of asparagine synthetase dictate the outcome of T-cell differentiation. **(A)** Box and whisker plot of qPCR-based time-course profile of *Asns* gene expression for CD8^+^ T cells FACS-sorted from mouse spleens during an *in vivo* primary immune response to LCMV infection. The *x* axis indicates the days post-infection on which cellswereanalyzed and the corresponding cell subpopulation identities, while the *y* axis represents gene expression values normalized relative to the average expression in the naïve state (d0). For those days where more than one cell subpopulation was sorted (d8 and d28), these are included as separate *x*-axis entries. The box hinges indicate the first/third quartiles of the data, while the mid-line represents the median, and the whiskers span the range of the data not considering outliers. Individual data points, each corresponding to an independent biological replicate, are indicated by the dots. Outliers (data points more than 1.5 x IQR away from the first/third quartiles, where IQR = inter-quartile range) are indicated in red. Statistical analysis based on one-way ANOVA to detect significant differences among the means of all subpopulations (p<0.0001), followed by Tukey’s multiple comparison tests between subpopulation pairs, with select p-values indicated in the plot. **(B)** FACS plots of CD25 *vs*. CD62L profiles for effector (IL-2)-polarized (**top**) and memory (IL-15)-polarized (**bottom**) cells at 120h differentiation, for cells activated and differentiated *in vitro* in asparagine-replete conditions (**left**), or maintained under conditions of asparagine depletion starting from 24h activation (**center**) or 0h differentiation (**right**). **(C)** FACS-based time-course profiles of fractional abundances of central-memory precursor (*Tcmp*, CD25^Lo^CD62L^Hi^) cells over 120h of effector (IL-2) polarization, for CD8^+^T cells activated and differentiated *in vitro* in asparagine-replete conditions (dark brown), or maintained under conditions of asparagine depletion starting from 24h activation (light brown), 0h differentiation (yellow), or at different time points during differentiation (dark to light blue). Statistical analysis based on F-Tests on 3^rd^ degree polynomials (see Methods), with global F-Test p-value shown on the bottom-right corner, and pairwise F-Test significance levels relative to the asparagine-replete condition displayed next to the legend entries for the different conditions. **(D)** FACS-based time-course profiles of fractional abundances of central-memory (*Tcm*, CD25^Lo^CD62L^Hi^) cells over 120h of memory (IL-15) polarization, for CD8^+^ T cells activated and differentiated *in vitro* in asparagine-replete conditions (dark green), or maintained under conditions of asparagine depletion starting from 24h activation (medium green), 0h differentiation (light green), or at different time points during differentiation (dark to light purple). Statistical analysis analogous to **(C)**. **(E-F)** FACS-based time-course profiles of fractional abundances of central-memory precursor **(E)** or central-memory **(F)** cells, for CD8^+^ T cells activated and differentiated *in vitro* in conditions analogous to those in **(C)** and **(D)**, but on a second experiment with slower differentiation kinetics, spanning up to 192h of differentiation, and with asparagine depletion performed only at 24h activation or 0h differentiation. Statistical analyses based on F-Tests on 4^th^ degree polynomials (see Methods), with global F-Test p-values shown on the top-left corners, and pairwise F-Test significance levels relative to the asparagine-replete condition displayed next to the legend entries for the different conditions. **(G-H)** qPCR-based time-course profiles of bulk *Asns* gene expression for effector (IL-2)-polarized (**G**) and memory (IL-15)-polarized (**H**) CD8^+^ T cells activated and differentiated *in vitro* in identical conditions as in **(E)** and **(F)**, and corresponding to the same experiment. The associated activation profiles (differentiation time < 0h) from the same experiment are included for completeness. The *y* axis represents expression values normalized relative to the average expression in the naïve state (−48h). Statistical analyses based on F-Tests on 5^th^ degree polynomials (see Methods), after log-transforming normalized expression levels X as log_2_(1+X), with global F-Test p-values shown on the top-left corners, and pairwise F-Test significance levels relative to the asparagine-replete condition displayed next to the legend entries for the different conditions. Significance levels corresponding to fixed-time pairwise t-Tests between either asparagine-depleted condition and the asparagine-replete condition, at time points immediately following each depletion, and at the later differentiation time points (144/168/192h), are indicated by the color-coded symbols on top of the corresponding data points (only for significant comparisons). Significance levels between the expression levels in each asparagine-depleted condition 24h after the respective depletions (0h *vs*. 24h) are indicated by the black symbols. The black lines indicate the lack of significant differences between both asparagine-depleted conditions at the later differentiation time points, based on fixed-time pairwise t-Tests. **(I)** FACS-based time-course profiles of relative fractions (out of total transduced cells) of control (EV, empty vector) and ASNS-OE CD8^+^ T cells isolated frommousespleens overa 28-day *in vivo*primary response to LCMV infection. Statistical analysis based on F-Test on 2^nd^ degree polynomials (see Methods), with the F-Test p-value shown on the bottom-left corner. The data for d0 are given by the experimental design, and were considered for F-Test statistical analysis. Significance levels corresponding to fixed-time pairwise t-Tests between either condition are indicated by the black symbols. **(J-K)** FACS-based plots of fractional abundances of short-lived effector cells (*SLEC*, KLRG1^Hi^CD127^Lo^) **(J)** and memory-precursor effector cells (*MPEC*, KLRG1^Lo^CD127^Hi^) **(K)**, for control (EV, empty vector) and ASNS-OE CD8^+^ T cells isolated from mouse spleens on day 8 of an *in vivo* primary response to LCMV infection. Statistical analyses based on two-tailed unpaired t-Tests with Welch’s correction (unequal variance assumption). **(L-M)** FACS-based time-course profiles of fractional abundances of central-memory (*Tcm*, CD44^Hi^CD62L^Hi^) cells **(L)**, and joint fractional abundances of effector and effector-memory (*Teff/Tem*, CD44^Hi^CD62L^Lo^) cells **(M)**, for control (EV, empty vector) and ASNS-OE CD8^+^ T cells isolated from mouse spleens over a 28-day *in vivo* primary response to LCMV infection. Statistical analyses analogous to **(I)**. All *in vitro* data correspond tothe averages of at least 3 independent culturewells per condition and timepoint, with error bars representing standard deviations (±SD). All *in vitro* FACS plots gated on live, single CD8^+^ T cells, and corresponding to the merge of 3 replicates. All *in vivo* data correspond to the averages of 3 biological replicates per condition and time point, with individual replicates indicated by the dots, and error bars representing standard errors of the mean (±SEM), unless otherwise noted.

Interestingly, in agreement with recent reports (Hope et al., 2021), we had shown above that CD8^+^ T cells are able to activate normally when extracellular asparagine is depleted after early activation (**Figures 2D-E, S2D-G**), presumably by increasing ASNS expression in response to this depletion. We thus asked whether altering asparagine availability, and thus potentially ASNS expression levels, could impact the fate of differentiating CD8^+^ T cells. To address this question, we activated CD8^+^ T cells *in vitro*, starting from asparagine-replete conditions, and further differentiated them under IL-2 (effector polarization) or IL-15 (memory polarization), while irreversibly depleting asparagine at multiple time points during differentiation, as well as after 24h activation (**Figure S4D**). We observed that neither cell viability (**Figures S4E-F**) nor proliferation (**Figure S4G**) were noticeably compromised in differentiating cells lacking an extracellular asparagine supply, regardless of whether depletion took place after 24h activation or at any point during differentiation. Strikingly, however, we did observe a shift towards a central-memory phenotype for cells differentiated in the absence of asparagine when depletion occurred after 24h activation, but not during differentiation (**Figure 4B**). Intriguingly, asparagine depletion during differentiation on the contrary led to a delay in central-memory polarization (**Figures 4C-D**), driven by a transient reduction in the fraction of central-memory precursor (*Tcmp*, for IL-2 polarization; **Figure 4C**) or central-memory (*Tcm*, for IL-15 polarization; **Figure 4D**) cells, relative to the asparagine-replete condition, immediately following asparagine removal. This was apparent regardless of when asparagine was withdrawn, but particularly significant when depletion occurred at the start of differentiation (**Figures 4C-D**), and was linked to an increase in the fraction of cells with an effector-like (*Teff*) phenotype under IL-2 polarization (**Figure S4H**). This phenotypic switch was further confirmed in independent experiments with slower differentiation kinetics (**Figures 4E-F, S4I**). In summary, we observe a markedly opposed response of CD8^+^ T cells to asparagine depletion during late activation or early differentiation *in vitro*, with the former promoting central-memory polarization, and the latter delaying it and favoring the maintenance of an effector phenotype.

We thus reasoned that, rather than asparagine availability, it is in fact the timing of asparagine depletion, and thus the potential dynamic modulation of ASNS expression, that may influence the outcome of CD8^+^ T-cell differentiation. To explore this hypothesis, we performed *in vitro* time-resolved measurements of *Asns* gene-expression for CD8^+^ T cells activated and further differentiated under IL-2 or IL-15, either in asparagine-replete conditions, or upon irreversible asparagine depletion at the start of differentiation or after 24h activation (**Figures 4G-H, S4D**). We observed a significant, transient increase in *Asns* expression following asparagine withdrawal both during late activation and early differentiation (**Figures 4G-H**). This supports the idea that, following early activation, CD8^+^ T cells can cope with a lack of extracellular asparagine by upregulating asparagine synthesis via ASNS (Hope et al., 2021). More importantly, however, our measurements show that, aside from this transient upregulation occurring earlier upon asparagine depletion during late activation, there are otherwise no differences between the expression profiles of cells lacking asparagine from late activation or early differentiation (**Figures 4G-H**). We also found no differences in *Asns* expression, in the long run, between the asparagine-depleted and asparagine-replete conditions under IL-2 polarization (**Figure 4G**), whereas these were rather minor under IL-15 polarization (**Figure 4H**). Taken together, these data support our hypothesis that it is the timing and dynamics of ASNS expression, rather than the extracellular availability of asparagine itself, which dictate the fate of CD8^+^ T-cell differentiation, with an earlier or later ASNS upregulation (and decay) respectively promoting or delaying the progression to a central-memory phenotype.

To further validate this hypothesis in a more physiological context, we next decided to disrupt the expression dynamics of ASNS by stably overexpressing it in CD8^+^ T cells differentiating *in vivo* in response to a viral infection. Specifically, we isolated naïve CD8^+^ T cells from P14 transgenic mice, and activated them *in vitro* for 24h in asparagine-replete conditions, followed by transduction with either control or ASNS-overexpression (ASNS-OE) retroviruses (**Figure S4J**). After an extra 24 hours of stimulation, we washed the transduced P14 cells, and co-transferred a 50:50 mixture of control and ASNS-OE cells into wild type C57BL/6J mice infected with LCMV Armstrong strain. We then evaluated their relative numbers and phenotypes at different time points during the ensuing immune response (**Figure S4J**). In line with our hypothesis that the dynamics of ASNS expression modulate CD8^+^ T-cell progression through the differentiation cascade, we found that preventing the decay of ASNS by stable overexpression *in vivo* led to a significant increase in the initial CD8^+^ T-cell expansion (**Figure 4I**). This was further accompanied by an apparent increase in the fraction of short-lived effector cells (**Figure 4J**), and a decrease in memory precursor frequency (**Figure 4K**) during the early stages of the immune response. Consistently with this promotion of an effector-like phenotype, we found that stable ASNS overexpression *in vivo* led to a significant delay in central-memory polarization (**Figure 4L**). This resulted in a significant decrease in the frequency of central-memory cells at the late stages of the response (**Figure 4L**), concomitant with an increase in the frequency of cells with an effector or effector-memory phenotype (**Figure 4M**). In conclusion, we find that disrupting the transient dynamics of ASNS expression during CD8^+^ T-cell differentiation *in vivo* promotes effector T-cell expansion, while simultaneously hindering long-term T-cell polarization towards a central-memory phenotype. Taken together, our *in vitro* and *in vivo* data thus demonstrate that the dynamics of ASNS expression dictate the fate of CD8^+^ T-cell differentiation.

## DISCUSSION

The metabolic rewiring undergone by CD8^+^ T cells transitioning along the immune response cascade is tightly linked to their functionality (Loftus and Finlay, 2016; Pearce and Pearce, 2017; Wang and Green, 2012). Yet, a comprehensive dynamic characterization of this metabolic rewiring is still missing (MacIver et al., 2013; Pearce et al., 2013). Here, we provide such dynamic description based on scRNA-seq measurements on CD8^+^ T cells progressing *in vitro* through the activation and differentiation cascade. Notably, our use of physiological nutrient conditions allows our scRNA-seq approach to capture a wide range of known aspects of *in vivo* T-cell metabolism, including particular features which were not recapitulated previously by *in vitro* models relying on commercial media (Levine et al., 2021; Ma et al., 2019). More importantly, our scRNA-seq approach allows us to further define new dynamic metabolic dependencies of activating and differentiating CD8^+^ T cells. Among them, we identify a critical role of the dynamics in asparagine synthetase (ASNS) expression in modulating CD8^+^ T-cell progression from an effector to a central-memory phenotype, which we further validate in an *in vivo* model of infection.

Despite the fact that the metabolism of T cells is known to evolve dynamically during an immune response, few studies to date have tried to characterize T-cell metabolism from a dynamic viewpoint. To date, these efforts were mostly based on elaborate time-resolved, bulk-level metabolic measurements, and were thus limited to the more tractable case of *in vitro* activating T cells (Edwards-Hicks et al., 2020; Wang et al., 2011). With the advent of single-cell omics, it is now not only possible to swiftly resolve the heterogeneity of cell states present in asynchronously-evolving cell populations (such as T cells progressing along the immune response cascade) (Grün, 2018; Stubbington et al., 2017), but also to simultaneously probe their metabolic states (Artyomov and Van den Bossche, 2020). As a result, a number of groups besides us have contributed to advancing our current understanding of the dynamics of T-cell metabolism by exploiting these single-cell approaches, particularly mass cytometry, both *in vitro* and *in vivo* (Ahl et al., 2020; Hartmann et al., 2021; Levine et al., 2021). Compared to scRNA-seq, mass cytometry provides a more direct readout of cellular metabolism, as it measures metabolic enzyme levels rather than gene expression, bypassing the potential impact of translation. Mass cytometry is also not subject to dropout effects as those observed in scRNA-seq, decreasing measurement variability among cells of a given state (Kharchenko et al., 2014). These advantages, however, come at the cost of coverage, given the lack of reliable antibodies for many metabolic enzymes (Rossi et al., 2021), and the limited number of proteins measurable in a single experiment (∼40, including the markers necessary for cell phenotyping) (Artyomov and Van den Bossche, 2020). Thus, scRNA-seq and mass cytometry should not be regarded as competing but rather as complementary approaches for studying metabolism at the single-cell level, with scRNA-seq providing a global way of identifying potentially interesting targets, whose metabolic impact may then be explored in a more direct and targeted way via mass cytometry.

Accordingly, and as shown by our data, thousands of genes can be simultaneously measured for every cell in an scRNA-seq experiment. Specifically, our identification of new dynamic metabolic dependencies of CD8^+^ T cells relied already on the joint exploration of over 1000 intracellular metabolic genes. Out of these, we focused on asparagine synthetase, but the potential impact of the expression dynamics of multiple other hits identified by our screen remains to be explored. Furthermore, our scRNA-seq data set includes also around 400 metabolite transporter genes, whose analysis we did not tackle, but which represent a potential route to provide additional insights on the dynamics of CD8^+^ T-cell metabolism. As shown by our correlation-based analysis of polyamine metabolism, scRNA-seq can capture the dynamic co-regulation between metabolic and signaling genes, as well as transcription factors. This approach may easily be extended to arbitrarily large gene networks, and coupled to regulatory-network identification algorithms (Aibar et al., 2017), to further investigate the dynamic link between transcriptional and metabolic programs in CD8^+^ T-cell responses. Thus, our work provides a validated, information rich resource for the immunometabolism community to further unravel new dynamic aspects of CD8+ T-cell metabolism. Furthermore, it provides a comprehensive and biologically well-defined data set that may be used as a model by the single-cell community to test and develop new methods for studying dynamic evolution processes.

Our identification of ASNS, and specifically of its expression dynamics, as a key modulator of CD8^+^ T-cell differentiation, is particularly timely. Indeed, several groups have recently reported critical roles for asparagine availability and ASNS expression in CD8^+^ T-cell activation (Hope et al., 2020; Torres et al., 2016; Wu et al., 2021). However, their potential impact on differentiation, and particularly memory formation, remained largely unexplored (Raynor and Chi, 2021). Our results thus complement those of these groups, by showing that, during CD8^+^ T-cell differentiation, it is not asparagine availability, but rather the dynamics of ASNS expression, that may impact both effector expansion and long-term memory formation. Although we did not delve into the mechanistic link between ASNS expression dynamics and effector expansion or memory formation, it is tempting to speculate that this may be tied to the regulation of mTORC1 activity. Indeed, like other amino acids, asparagine has been reported to activate mTORC1 (Krall et al., 2021; Meng et al., 2020). The dynamic modulation of mTORC1 activity, in turn, has been shown to determine both the naïve-to-effector and effector-to-memory transitions in CD8^+^ T cells (Rao et al., 2010). Thus, based on our *in vivo* data, one may speculate that early ASNS overexpression could lead to an early accumulation of intracellular asparagine, driving the observed increase in early effector CD8^+^ T-cell expansion via enhanced mTORC1 activation. On the other hand, preventing the later downregulation of ASNS expression would make these elevated intracellular asparagine levels persist, leading to a chronic mTORC1 overactivation, in turn hindering long-term memory formation. Of note, the link between ASNS expression and mTORC1 activity would not need to be exclusively mediated by asparagine itself. It may also be linked to an enhanced exchange of intracellular asparagine for other amino acids in the extracellular environment such as arginine, serine, or histidine (Krall et al., 2016), all of which are known mTORC1 activators (Meng et al., 2020).

Interestingly, despite the observed impairment in central-memory polarization upon ASNS overexpression, the latter may in fact represent an attractive approach for immunometabolic modulation in adoptive cell transfer immunotherapies, particularly against tumors susceptible to asparaginase treatment, as the latter has been shown to hinder T-cell responses (Wu et al., 2021). The rationale for this is that, despite the fractional decrease we observe in the central-memory compartment upon ASNS overexpression, the absolute numbers of central-memory cells are actually doubled relative to the control at the late stages of the response, due to the enhanced initial effector expansion (**Figure S4K**). Thus, one may speculate on how transient rather than stable ASNS overexpression strategies, based on genetic approaches (Kim and Eberwine, 2010), or perhaps on asparagine depletion during late *ex vivo* activation prior to transfer, may be of interest for further studies. Indeed, one may expect for these approaches to maintain the enhanced initial effector expansion, while avoiding the deleterious impact on long-term memory formation.

In conclusion, we provide a resource on the dynamic metabolic changes undergone by CD8^+^ T cells when transitioning along the immune response cascade. We expect that this resource will provide the basis for identifying novel strategies to modulate T-cell functionality.

## ACKNOWLEDGEMENTS

We are grateful to Stephanie Humblet-Baron, Adrian Liston, and Massimiliano Mazzone for guidance on the immunology aspects of the study and for providing reagents. We also thank Junbin Qian, Bram Boeckx, Florian Rambow, Bernard Thienpont, and Asaf Madi for help and feedback on scRNA-seq data analysis and interpretation. Finally, we are also grateful to all members of the Fendt lab for their valuable suggestions and feedback on the manuscript. Parts of Figures S1A, S2C, and S4C/D/J were created with BioRender.com.

JF-G was supported by consecutive FWO Junior and Senior postdoctoral fellowships. S-MF acknowledges funding from the European Research Council under the ERC Consolidator Grant Agreement n. 771486–MetaRegulation, FWO–Research Projects and Fonds Baillet Latour. P-CH is funded in part by the European Research Council Starting Grant (802773-MitoGuide), the SNSF project grants (31003A_182470), the Cancer Research Institute (CLIP investigator award and Lloyd J. Old STAR award), University of Lausanne, and Ludwig Cancer Research. PC is supported by Grants from Methusalem funding (Flemish government), the Fund for Scientific Research-Flanders (FWO-Vlaanderen), ERC Advanced Research Grant EU-(ERC743074), and an NNF Laureate Research Grant from Novo Nordisk Foundation (Denmark).

---

**Figure S1.**
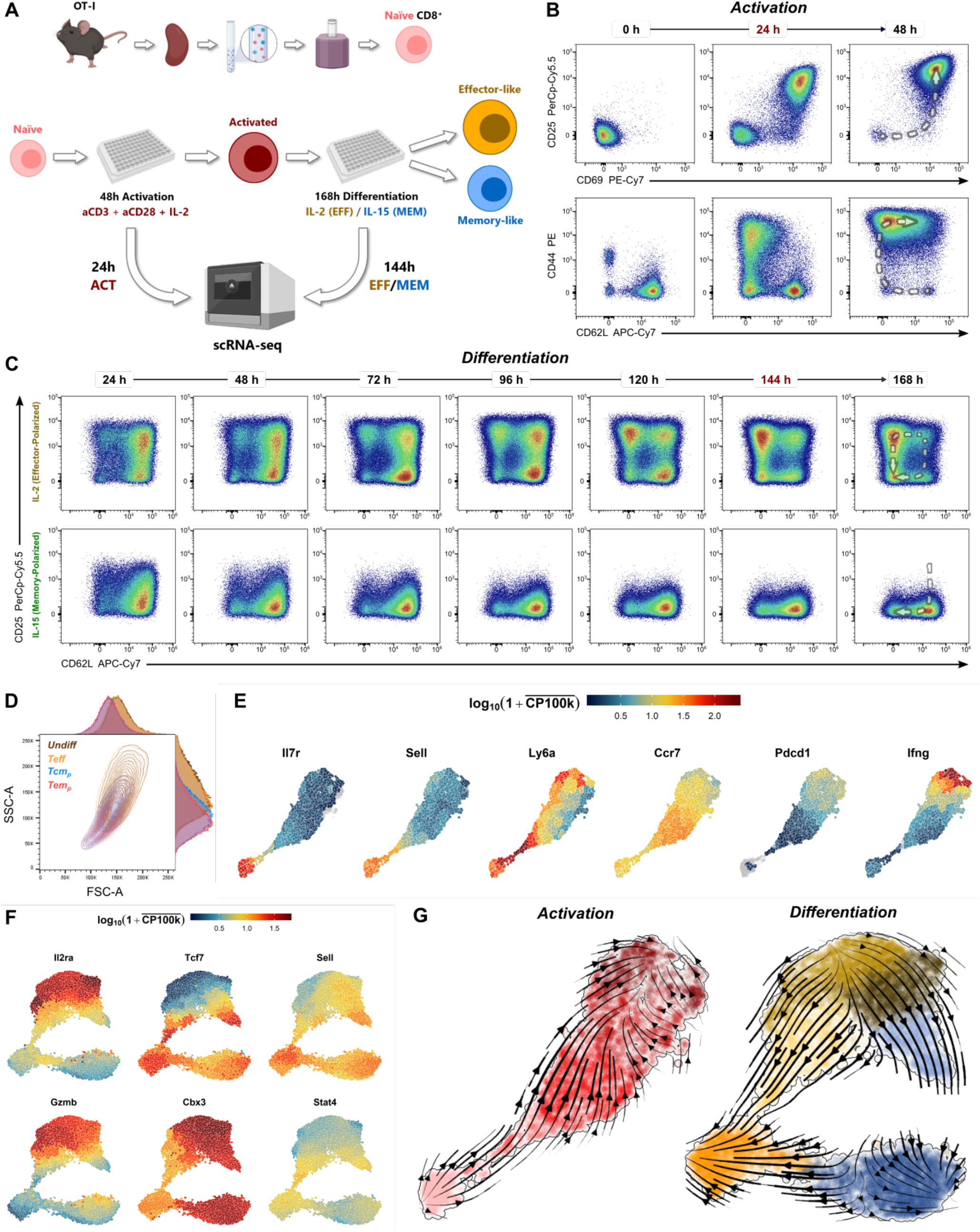
Single-cell RNA-seq captures the population dynamics of *in vitro* activating/differentiating CD8^+^ T cells. **(A)** Schematic of the experimental approach used to perform scRNA-seq measurements on *in vitro* activating/differentiating CD8^+^ T cells. **(B–C)** Time-resolved FACS plots of CD25 *vs*. CD69 (**top**) and CD44 *vs*. CD62L (**bottom**) profiles along 48h of *in vitro* CD8^+^ T-cell activation **(B)**, and of CD25 *vs*. CD62L profiles along 168h of subsequent *in vitro* effector (IL-2) (**top**) and memory (IL-15) polarization (**bottom**). The dashed arrows over the endpoints (48h activation and 168h effector/memory polarization) indicate the dynamic evolution of the various surface markers along the activation/differentiation process. The optimal time points used for scRNA-seq are indicated in red. **(D)** FACS plot of FSC-A *vs*. SSC-A, and corresponding histogram plots, for the 4 different populations identified in our 144h effector-polarized sample, showing the size and morphology differences between the *Undiff/Teff* and *Tcm*_*p*_/Tem_p_ populations. **(E-F)** UMAP plots for the scRNA-seq data corresponding to the 24h activation sample **(E)** and the two 144h differentiation samples combined **(F)**, color-coded based on the expression levels of the key marker genes used for cell-state annotation. Log-transformed gene expression levels are indicated by the color scale, where 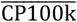 denotes the average gene expression level (in counts per 100k reads) over all cells in a given fine-grained cluster (see Methods), and with the cells with highest expression levels plotted on top. The *Gzmb* expression levels in **(F)** were scaled down by a factor of 10 before log-transformation, to avoid distorting the color code for the remaining genes, and therefore the color-code represents 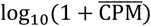 for *Gzmb*, where 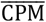 denotes counts per million reads. **(G)** RNA velocity stream plots for the scRNA-seq data corresponding to the 24h activation sample **(left)** and the two 144h differentiation samples combined **(right)**. The underlying UMAP plots and color-coding scheme are identical to those in Figures 1F-G.

**Figure S2.**
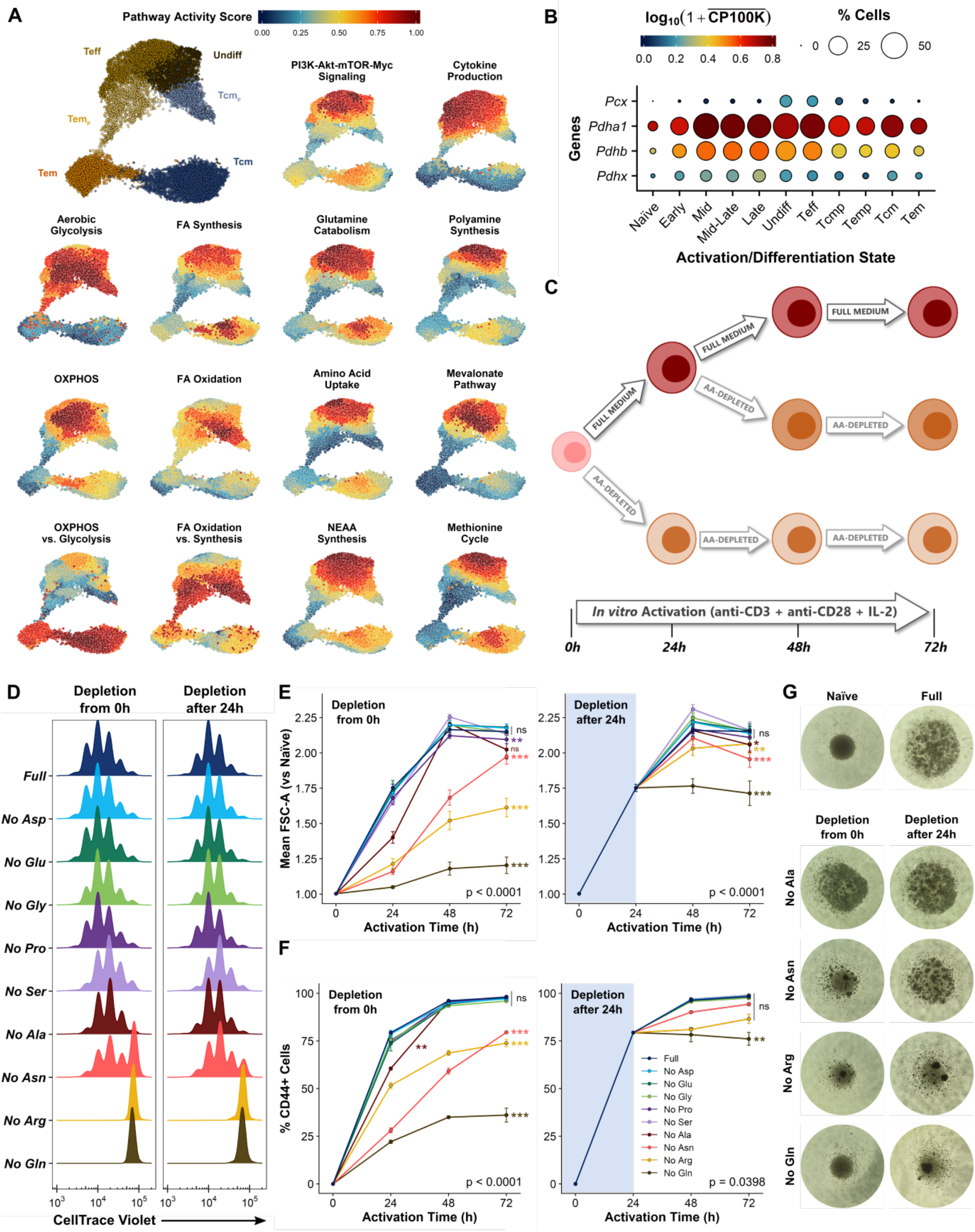
Single-cell RNA-seq identifies known aspects of *in vivo* CD8^+^ T-cell metabolism during activation and differentiation and predicts the dependence of activating CD8^+^ T cells on amino acid uptake *vs*. synthesis. **(A)** UMAP plots for the scRNA-seq data corresponding to the two 144h differentiation samples combined, color-coded based on GSVA-based pathway activity scores for identical metabolic and/or signaling pathways as in **Figure 2A**. Pathway activity scores are averaged over all cells in a given fine-grained cluster, and further scaled to the range 0–1 for each individual pathway (see Methods). A replica of Figure 1G is included for reference. **(B)** Gene expression *vs*. cell state profiles for select genes involved in pyruvate metabolism. Log-transformed gene expression levels are indicated by the color scale, where 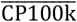 denotes the average gene expression level (in counts per 100k reads) over all cells in a given cell state, while the areas of the circles represent the percentage of cells with non-zero expression of each gene among all cells in each cell state. **(C)** Schematic depiction of our experimental approach for studying the impact of non-essential amino acid depletion on the *in vitro* activation of CD8^+^ T cells. **(D)** CellTrace^®^ Violet dilution profiles after 72h of *in vitro* stimulation in nutrient-replete BLM (Full), or upon single-NEAA depletion, either from the start of the stimulation **(left)**, or after 24h stimulation in nutrient-replete BLM **(right)**. Data correspond to averages of 4 independent culture wells per condition, and are representative of 2 independent experiments. **(E-F)** FACS-based time-course profiles for the average cell-size (mean FSC-A intensity) **(D)** and extent of activation (percentage of CD44^Hi^ cells) **(E)** over a total of 72h of stimulation, for identical NEAA depletion conditions as in **(D)**. Data correspond to averages of 4 independent culture wells per condition and time point, and are representative of 2 independent experiments. Error bars represent standard deviations (±SD). Statistical analyses based on F-Tests on 2^nd^ degree polynomials (see Methods), with global F-Test p-values shown on the bottom-right corners, and pairwise F-Test significance levels relative to the Full condition displayed next to the profiles for the different conditions. **(G)** Representative photographs of CD8^+^ T cells in the absence of stimulation (Naïve), or stimulated for 48h in nutrient-replete BLM (Full), or upon depletion of select NEAAs, either from the start of the stimulation **(left)**, or after 24h stimulation in nutrient-replete BLM **(right)**.

**Figure S3.**
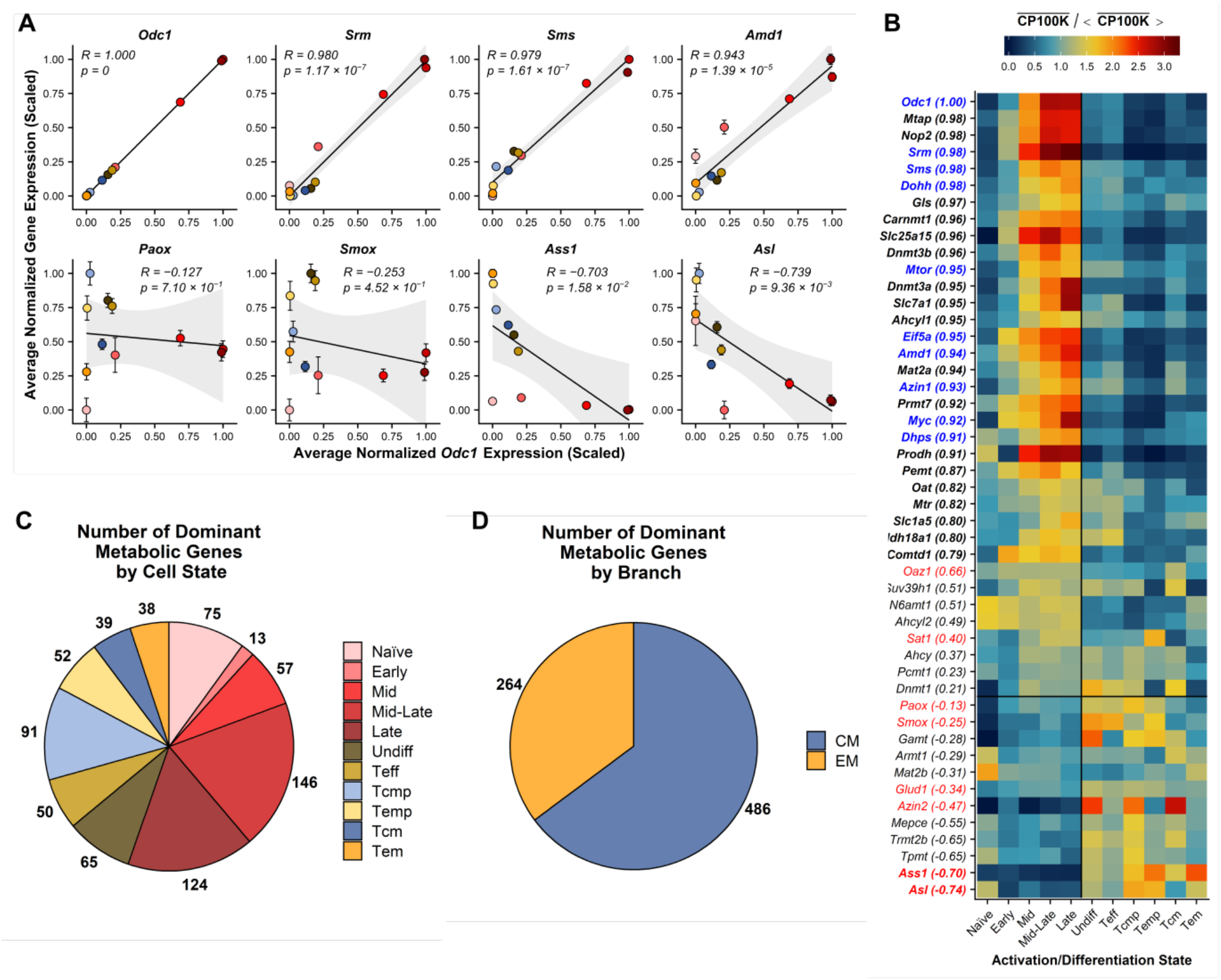
Single-cell RNA-seq captures the dynamic orchestration of CD8^+^ T-cell metabolic programs during activation and differentiation and identifies novel dynamic metabolic dependencies. **(A)**Correlation plots between the expression profiles of the various genes shown in **Figure 2A** and that of *Odc1*. The *y* values represent the cell state-averaged expression levels for each of the corresponding genes in every cell state, while the *x* values represent in all cases those for *Odc1* (see Methods). The corresponding Pearson correlation coefficient (*R*) values and associated p-values are displayed in each plot, with solid lines representing the best linear fits of the data, and the grey shaded areas showing the 95% confidence intervals for the fits. The various activation/differentiation states are indicated via the dot color code, matching the color code in **Figure 2A**. Error bars represent standard errors of the mean (±SEM) for the cell state-averaged expression levels of each of the corresponding genes, considering the expression levels of all cells in each state. **(B)**Heatmap plot of cell state-averaged expression profiles for the polyamine metabolism genes shown in **Figure 2B**. Scaled expression levels are indicated by the color scale, where 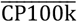 denotes the average gene expression level (in counts per 100k reads) over all cells in a given cell state, and 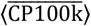 the average of the latter over all cell states. Genes are sorted by their Pearson-*R* values relative to *Odc1* (shown in parentheses by the gene names), with genes with |*R*| *> 0*.*7* indicated in bold face. Genes involved in polyamine synthesis and hypusination, or being known upstream regulators or downstream targets of the latter, are highlighted in blue, while genes opposing polyamine synthesis are highlighted in red. **(C-D)** Pie charts indicating the number of metabolic genes presenting a dominant expression in each activation/differentiation state **(C)**, or along the differentiation branches leading to the generation of either a *Tcm* (CM = *Undiff* + *Tcm*_*p*_ + *Tcm*) or a *Tem* (EM = *Teff* + *Tem*_*p*_ + *Tem*) phenotype **(D)**.

**Figure S4.**
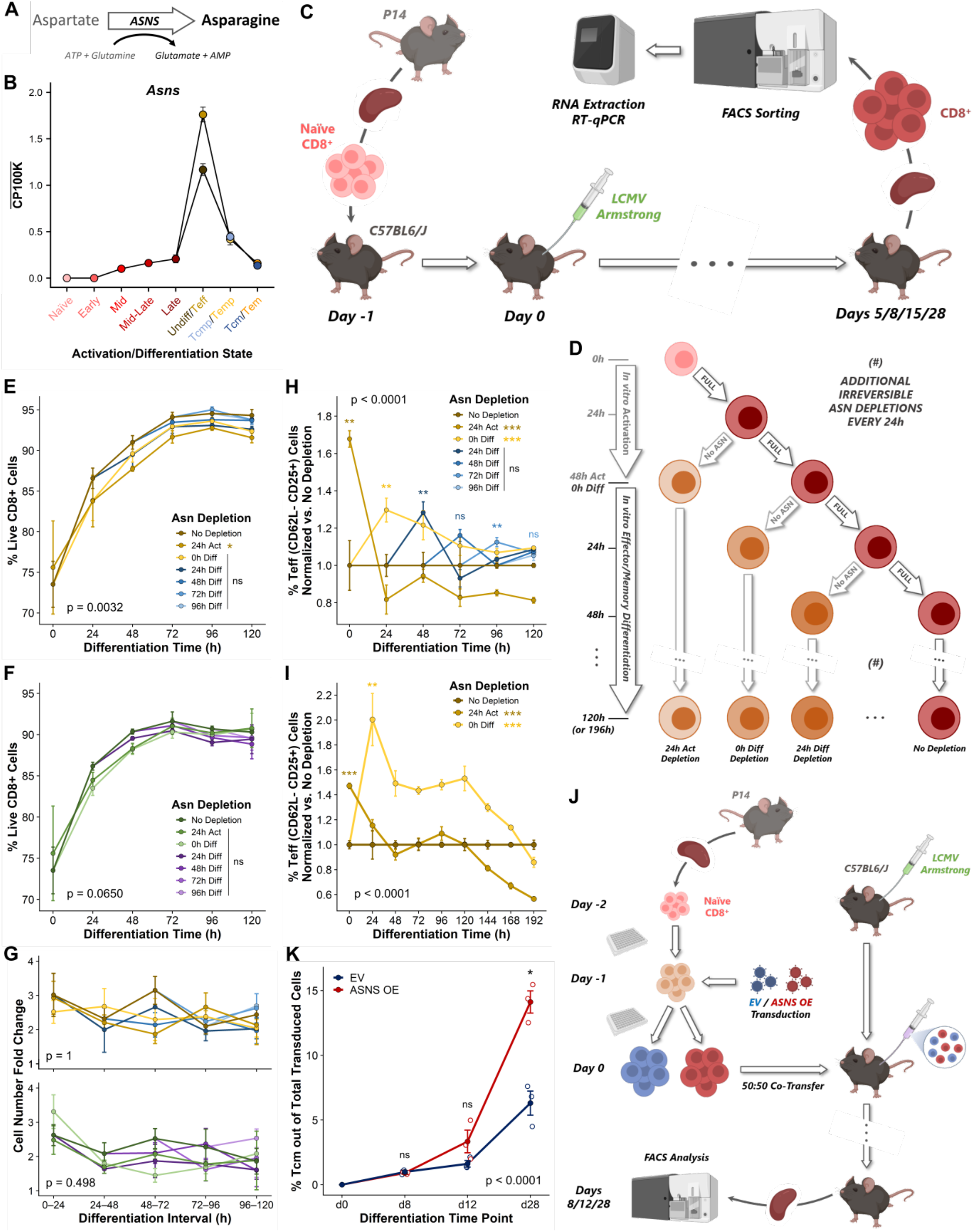
The expression dynamics of asparagine synthetase dictate the outcome of T-cell differentiation. **(A)**Schematic representation of the asparagine synthetase (ASNS) enzymatic reaction. **(B)**Cell state-averaged gene expression profile for asparagine synthetase (*Asns*), based on our full merged scRNA-seq data set. The *y*-axis values represent normalized *Asns* expression levels (in counts per 100k reads) averaged over all cells in each given activation/differentiation state, with cell states indicated both in the *x* axis and via the dot color code. Error bars represent standard errors of the mean (±SEM), considering the expression levels of all cells in each state. **(C)**Schematic depiction of our experimental approach for studying the dynamics of *Asns* gene expression in CD8^+^ T cells during an *in vivo* primary immune response to LCMV infection. **(D)**Schematic depiction of our experimental approach for studying the impact of asparagine depletion on the *in vitro* differentiation of CD8^+^ T cells. **(E)**FACS-based time-course profiles of total viable cell fractions over 120h of effector (IL-2) polarization, for CD8^+^ T cells activated and differentiated *in vitro* in asparagine-replete conditions (dark brown), or maintained under conditions of asparagine depletion starting from 24h activation (light brown), 0h differentiation (yellow), or at different time points during differentiation (dark to light blue). The data correspond to the same experiment as **Figure 4C**. Statistical analysis based on F-Tests on 3^rd^ degree polynomials (see Methods), with global F-Test p-value shown on the bottom-left corner, and pairwise F-Test significance levels relative to the asparagine-replete condition displayed next to the legend entries for the different conditions. **(F)**FACS-based time-course profiles of total viable cell fractions over 120h of memory (IL-15) polarization, for CD8^+^ T cells activated and differentiated *in vitro* in asparagine-replete conditions (dark green), or maintained under conditions of asparagine depletion starting from 24h activation (medium green), 0h differentiation (light green), or at different time points during differentiation (dark to light purple). The data correspond to the same experiment as **Figure 4D**. Statistical analysis analogous to **(E)**. **(G)**Time-course profiles of daily cell-number fold-changes (based on cell counts) over 120h of effector (IL-2) **(top)** or memory (IL-15) **(bottom)** polarization, for CD8^+^ T cells activated and differentiated *in vitro* in identical conditions as in **(E)** and **(F)**, and corresponding to the same experiments. Statistical analyses based on F-Tests on 3^rd^ degree polynomials (see Methods), with global F-Test p-values shown on the bottom-left corners (both not significant). **(H)**FACS-based time-course profiles of fractional abundances of effector (*Teff*, CD25^Hi^CD62^Lo^) cells over 120h of effector (IL-2) polarization, for CD8^+^ T cells activated and differentiated *in vitro* under conditions of asparagine depletion starting from 24h activation (light brown), 0h differentiation (yellow), or at different time points during differentiation (dark to light blue), normalized relative to the corresponding *Teff* fractional abundances for cells activated and differentiated in asparagine-replete conditions (dark brown). The data correspond to the same experiment as **Figure 4C**. Statistical analysis based on F-Tests on 3^rd^ degree polynomials (see Methods), with global F-Test p-value shown on the top-left corner, and pairwise F-Test significance levels relative to the asparagine-replete condition displayed next to the legend entries for the different conditions. Significance levels corresponding to fixed-time pairwise t-Tests between either asparagine-depleted condition and the asparagine-replete condition, at time points immediately following each depletion, are indicated by the color-coded symbols on top of the corresponding data points. **(I)**FACS-based time-course profiles of fractional abundances of effector (*Teff*, CD25^Hi^CD62^Lo^) cells over 192h of effector (IL-2) polarization, for CD8^+^ T cells activated and differentiated *in vitro* under conditions of asparagine depletion starting from 24h activation (light brown) or 0h differentiation (yellow), normalized relative to the corresponding *Teff* fractional abundances for cells activated and differentiated in asparagine-replete conditions (dark brown). The data correspond to the same experiment as **Figure 4E**. Statistical analysis analogous to **(I)**, but using 4^th^ degree polynomials. **(J)**Schematic depiction of our experimental approach for studying the impact of stable ASNS overexpression on the differentiation of CD8^+^ T cells during an *in vivo* primary immune response to LCMV infection. **(K)**FACS-based time-course profiles of relative fractions (out of total transduced cells) of central-memory (*Tcm*, CD44^Hi^CD62L^Hi^) cells, for control (EV, empty vector) and ASNS-OE CD8^+^ T cells isolated from mouse spleens over a 28-day *in vivo* primary response to LCMV infection. The data originate from multiplying those in Figures 4I and 4L. Statistical analysis based on F-Test on 2^nd^ degree polynomials (see Methods), with the F-Test p-value shown on the bottom-left corner. The data for d0 are given by the experimental design, and were considered for F-Test statistical analysis. Significance levels corresponding to fixed-time pairwise t-Tests between either condition are indicated by the black symbols. All *in vitro* data except for the scRNA-seq data in **(B)** correspond to the averages of at least 3 independent culture wells per condition and time point, with error bars representing standard deviations (±SD). All *in vitro* FACS plots gated on live, single CD8^+^ T cells, and corresponding to the merge of 3 replicates. All *in vivo* data correspond to the averages of 3 biological replicates per condition and time point, with individual replicates indicated by the dots, and error bars representing standard errors of the mean (±SEM), unless otherwise noted.

## METHODS

### Mice

For *in vitro* experiments, OT-I transgenic (C57BL/6-Tg(TcraTcrb)1100Mjb/Crl) mice, with a T-cell receptor designed to recognize ovalbumin peptide residues 257–264 (OVA_SIINFEKL_) (Clarke et al., 2000; Hogquist et al., 1994), were obtained from Charles River (France), and were housed in individually ventilated cages within specific pathogen-free (SPF) facilities at the KU Leuven (Leuven, Belgium) until the start of experiments. All animal experiments and experimental procedures were previously approved by the KU Leuven Ethical Committee for Animal Experimentation (ECD, Project No. P158-2016), and were performed in compliance with all relevant ethical regulations and adhering to the triple-R principle (replacement/reduction/refinement).

For *in vivo* experiments, P14 transgenic mice (B6.Cg-Tcra^tm1Mom^Tg(TcrLCMV)327Sdz/TacMmjax), with a T-cell receptor designed to recognize peptide gp33 (KAVYNFATM) from lymphocytic choriomeningitis virus (LCMV) (Mombaerts et al., 1992; Pircher et al., 1989), were obtained from The Jackson Laboratory (Bar Harbor, ME, USA). Wild type C57BL/6J mice were obtained from The Jackson Laboratory and bred in house. Animals were housed in individually ventilated cages within SPF facilities at the University of Lausanne (Lausanne, Switzerland). All animal experiments and experimental procedures were previously approved and performed in accordance with the guidelines and regulations implemented by the Swiss Animal Welfare Ordinance.

### Mouse T-cell isolation for in vitro experiments

Naïve CD8^+^ T cells were isolated from the spleens of 8–13-week-old OT-I transgenic mice. Mice were sacrificed with an overdose of a highly concentrated barbiturate (Dolethal^®^), injected intraperitoneally at a dosage of 150-200 mg/kg. A quick whole-body perfusion with cold PBS was then performed before harvesting the spleen. Splenocyte suspensions were prepared in Separation Buffer (SB: PBS + 3% FBS + 2 mM EDTA), by mashing the freshly-resected whole spleens through 70 µm nylon cell strainers (VWR) using the rubber ends of 10 mL syringe plungers (Terumo). Splenocytes were then resuspended in 4 mL/spleen ice-cold Red Blood Cell Lysis Buffer (Sigma-Aldrich) and incubated for 5 minutes on ice, after which they were washed twice with SB, filtered again through 70 µm nylon cell strainers, and resuspended at 10^8^ cells/mL in SB. Naïve CD8^+^ T cells were then negatively-selected (CD8^+^CD44^-^ purity > 90%) from the resulting single-cell splenocyte suspension using the MojoSort™ Mouse CD8 Naïve T Cell Isolation Kit (BioLegend, San Diego, CA, USA). Two modifications were introduced relative to the manufacturer’s guidelines to increase purity: antibody/bead incubations were carried out at room temperature (instead of on ice), and cells were gently resuspended halfway through the bead incubation. Isolated naïve CD8^+^ T cells were then either stained for proliferation tracking, or directly prepared for plating, as described below.

### CellTrace™ Violet in vitro proliferation-tracking assays

To allow tracking the proliferation of CD8^+^ T cells upon activation in particular *in vitro* experiments, freshly-isolated naïve CD8^+^ T cells were resuspended at a density of ∼10^7^ cells/mL in pre-warmed (37°C) SB supplemented with 5 µM CellTrace™ Violet reagent (Invitrogen), and gently dispersed by pipetting. Cell suspensions were then incubated for 12 minutes in the dark inside a water bath at 37°C, and gently shaken every 3 minutes to prevent aggregation. After that, cells were washed with 10 volumes of room-temperature SB, to neutralize and remove the excess CellTrace™ dye, and prepared for plating, as described below.

### In vitro CD8^+^ T-cell activation and effector/memory polarization

Flat- or round-bottom 96-well plates (Corning) were coated overnight at 4°C with 50 µL/well of either 10 µg/mL (flat-bottom) or 5 µg/mL (round-bottom) anti-mouse CD3ε (BioLegend) in PBS (for stimulated wells), or 50 µL/well of PBS (for unstimulated controls). Prior to cell seeding, plates were washed twice with 200 µL/well PBS. Freshly isolated naïve CD8^+^ T cells were then seeded at ∼0.5×10^6^ cells/mL (200 µL/well = 10^5^ cells/well) in culture medium supplemented with 0.5 µg/mL soluble anti-mouse CD28 (eBioscience) and 10 ng/mL (∼50 U/mL) recombinant murine Interleukin-2 (IL-2, BioLegend), and activated for up to 72h. For differentiation experiments, cells were harvested after 50-54h activation, counted, and re-plated in new round-bottom 96-well plates at ∼0.25×10^6^ cells/mL (200 µL/well = 5×10^4^ cells/well) in fresh culture medium supplemented with either 10 ng/mL IL-2, for effector polarization, or 50 ng/mL (∼10 U/mL) recombinant murine Interleukin-15 (IL-15, Peprotech), for memory polarization. Cells were then differentiated for up to 192h in either condition, subject to daily harvesting, counting, and re-plating in new round-bottom 96-well plates at ∼0.5×10^6^ cells/mL (200 µL/well = 10^5^ cells/well) in fresh culture media supplemented with either cytokine (IL-2 for effector polarization, or IL-15 for memory polarization). In experiments involving nutrient depletion during differentiation, or at the beginning of activation/differentiation, this was achieved by simply resuspending the cells in the appropriate nutrient-depleted (rather than nutrient-replete) culture medium prior to seeding/re-plating. In experiments involving nutrient depletion from the culture media after 24h activation, the plated cells were first spun down (3 minutes at 300 x g, room temperature) and medium was aspirated from each well. Cells were then washed once with fresh medium lacking the depleted nutrients, followed by a second spin-down and medium aspiration, and final addition of nutrient-depleted medium supplemented with anti-CD28 and IL-2 (same concentrations as above). Media aspirations and additions were performed gently and along the edges of the plate wells, to minimize perturbations on the activating cells. Nutrient-replete controls were subject to an identical medium replacement approach as nutrient-depleted conditions, only adding nutrient-replete medium in the final step. Round-bottom plates were used for activation in all experiments involving nutrient depletion (since this minimizes cell losses and perturbations during medium aspirations), whereas flat-bottom plates were used for activation in all other experiments (specifically, for scRNA-seq measurements). The respective anti-CD3ε concentrations were selected accordingly, to achieve identical activation dynamics in both cases.

### In vitro culture conditions

All *in vitro* experiments were performed in humidified temperature/CO_2_-controlled (37°C, 5% CO_2_) incubators, and using a custom home-made blood-like medium (BLM), whose complete formulation and preparation has been described elsewhere (Fernández-García and Fendt, 2019). BLM was in all cases supplemented with 10% fetal bovine serum (FBS), 100 U/mL penicillin, 100 µg/mL streptomycin, and 50 µM β-mercaptoethanol (all from Gibco). In experiments involving nutrient depletion, FBS was replaced with dialyzed FBS, to ensure full nutrient depletion, and the pH of nutrient-depleted or replete BLM stocks was adjusted individually for each formulation to achieve identical pH conditions in all of them. IL-2, IL-15, and anti-CD28 were always added freshly to the culture media, immediately before cell resuspension, and serum-containing BLM aliquots were always used within 3 days of preparation, to mitigate serum-driven nutrient (e.g. glutamine) degradation.

### Dead cell removal and single-cell RNA sequencing cell preparation

T-cell activation is necessarily accompanied by substantial cell death. In order to mitigate the impact of the presence of dead cells in our downstream single-cell sequencing measurements, dead cells were removed from our 24h activation sample (∼50% viability) using a MACS^®^ Dead Cell Removal Kit (Miltenyi Biotec). In brief, ∼2×10^6^ cells were pooled from 24 culture wells and resuspended in 100 µL Dead Cell Removal MicroBeads suspension, following the manufacturer’s guidelines thereafter. The purified live-cell suspension (∼90% viability) was then washed once with fresh culture medium, and resuspended at ∼10^6^ cells/mL in fresh culture medium supplemented with 10 ng/mL IL-2. Dead cell removal was not necessary for our 144h effector-polarized (∼90% viability) or memory-polarized (∼70% viability) samples, in which case ∼10^6^ cells/sample were pooled from 6 culture wells each, washed once with fresh culture media, and resuspended at ∼10^6^ cells/mL in fresh culture media supplemented with either 10 ng/mL IL-2 (for the effector-polarized sample) or 50 ng/mL IL-15 (for the memory-polarized sample). Cell suspensions were kept on ice and immediately processed for single-cell library preparation, with a fraction of each cell suspension preserved for simultaneous FACS analysis. The choice of culture medium for cell resuspension before single-cell library preparation, rather than the standard PBS + 0.04% BSA buffer, was motivated by the fact that preliminary tests with the latter resulted in a noticeable decrease in cell quality, likely due to cell starvation resulting from the elevated metabolic activity of CD8^+^ T cells. This choice was further motivated by the focus of our study in quantifying metabolic gene expression, for which maintaining cells in a nutrient-depleted environment would most likely lead to spurious results.

### Single-cell RNA sequencing, data pre-processing, and cell-state assignment

Cell suspensions for each sample were converted to barcoded single-cell cDNA libraries using the Chromium Single Cell 3’ V2 Library, Gel Bead, Chip, and Multiplex Kit (10x Genomics), following the manufacturer’s guidelines, and aiming for a total of 10000 cells per library. Single-cell libraries were then sequenced on a NovaSeq 6000 System (Illumina). The sequenced reads were then mapped to the mouse genome (mm10 build GRCm38.p4) using the Cell Ranger software (10x Genomics), and the resulting single-cell gene expression data was analyzed within the *R/Bioconductor* framework. Specifically, the raw UMI count matrices for each individual sample were first imported using *Seurat* (Butler et al., 2018), and then converted for further processing with *Monocle* (Cao et al., 2019). Low-quality cells were then filtered based on standard quality-control metrics, with sample-specific thresholds chosen based on evaluating quality-control histograms for each sample independently. In particular, cells were filtered based on their mitochondrial RNA content (allowing for a maximum of 5% in all cases), library size (removing cells with total UMI counts below 1000/2500/2000 for the activation/effector/memory samples, respectively), and number of detected genes (removing cells expressing less than 350/800/700 genes for the activation/effector/memory samples, respectively). Genes expressed in less than 10 cells were additionally ignored in all subsequent analyses. Size-factor and variance-stabilizing normalization (based on fitting to a negative binomial distribution) were then applied to the filtered data sets, and highly-variable genes (HVGs) were identified for each of them based on their departure from the average normalized dispersion *versus* expression trend observed among all genes. After excluding mitochondrial, ribosomal-protein, and cell cycle-associated genes, the top 1000 HVGs with size-factor normalized expressions above 0.005 were selected. Principal component analysis (PCA) was then performed on the size factor-normalized and variance-stabilized count matrix restricted to these genes only, followed by 2D UMAP dimensional reduction (McInnes et al., 2018) based on the resulting top 50 principal components (with *correlation* distance metric, *number of neighbors* = 15, and *minimum distance* = 0.05, and without further PCA scaling). After that, cells were clustered in the UMAP plane by applying the *Louvain* (Blondel et al., 2008) graph-based algorithm at high resolution (*k*_*NN*_ = 5/10/5 for the activation/effector/memory samples, respectively), in order to attain a fine-grained cluster structure for each sample (69/66/40 clusters for the activation/effector/memory samples, respectively). The resulting fine-grained clusters were then manually annotated to specific cell states, based on evaluating the cluster-averaged normalized expression profiles of a number of cell state-specific markers. Cells clearly clustering away from the bulk of the cells in each sample in the UMAP space, or clusters displaying features indicative of low quality (e.g. elevated *Actb* content, or low number of detected genes), were further removed for all subsequent analyses at this point.

Merged samples, consisting either of both the effector- and memory-polarized samples (differentiation), or all 3 samples put together (full) were processed following an analogous approach to the one described above for the single samples, with some particularities. First, cell filtering was performed simply by removing all cells not present in any of the final single-sample data sets. Second, UMAP dimensional reduction was performed using *cosine* distance metric and increased *number of neighbors* (30/65 for the differentiation/full merged samples, respectively) and *minimum distance* (0.2/0.4 for the differentiation/full merged samples, respectively) parameters, to account for the increased cell numbers. In the case of the full merged sample, batch regression was further included in the PCA step, to account for the fact that the cDNA libraries for the activation and differentiation samples were prepared on different days. Finally, formal clustering was not performed on the merged samples, but instead the cell-state annotations derived from the single samples were directly transferred to their merged counterparts.

### Trajectory inference and RNA velocity

Trajectory inference was performed on the combination of all 3 samples within the *Monocle3-alpha* framework (Cao et al., 2019), by applying reversed graph embedding in the UMAP plane based on the *SimplePPT* algorithm, using default parameters (specifically, without further gene expression scaling) other than forcing a single Louvain partition and allowing for closed-loop trajectories. RNA velocity calculations were performed separately on the activation sample and the combination of both differentiation samples (effector + memory), within the *scVelo* framework (Bergen et al., 2020). For this, the *BAM* files generated by the Cell Ranger software (10x Genomics) were first processed with *velocyto* (La Manno et al., 2018) (without applying a repeat annotation mask), in order to obtain *Loom* files with spliced/unspliced counts for each of the 3 samples. The *Loom* files for the two differentiation samples (effector + memory) were then combined using *loompy* (www.loompy.org), and *scVelo*-compatible *AnnData* (.h5ad) files for the filtered activation and differentiation (effector + memory) data sets were generated using the *R* package *sceasy* (www.github.com/cellgeni/sceasy). The corresponding *Loom* and *AnnData* files were then combined within *scVelo* (Bergen et al., 2020), and processed in order to obtain RNA velocity profiles. RNA velocity calculations were based on the top 50 principal components resulting from applying PCA to the top 2000 highly-variable genes in the data set (after filtering out genes with fewer than 20 total counts), with first/second-order moment computations for each cell taking into account the 500 nearest neighbors. The simpler, *stochastic* velocity model was used to determine RNA velocities for the differentiation (effector + memory) sample, whereas the more complex *dynamical* model was used for the activation sample. This choice was imposed by the fact that, during CD8^+^ T-cell activation, a majority of the top highly-variable genes corresponds to genes whose expression is repressed along activation, leading to the *stochastic* model yielding velocities inverted relative to the expected biological progression.

### Pathway activity analysis

Pathway activity analysis was performed using the *GSVA* (gene set variation analysis) package (Hänzelmann et al., 2013). In brief, count matrices for the filtered activation and differentiation (effector + memory) data sets were first subject to size-factor and variance-stabilizing normalization. *GSVA* scores were then determined using default parameters (particularly a Gaussian kernel for cumulative density function estimation, as the input expression matrix is already log-normalized by virtue of the variance stabilization) for a manually assembled list of gene sets (see **Supplementary Table 2**), each including genes positively (UP) or negatively (DOWN) correlated with the activity of various metabolic and signaling pathways. Activity scores for each pathway were then determined for every cell by subtracting the *GSVA* scores determined for the corresponding UP and DOWN gene sets. Pathway activity scores were then averaged over all cells belonging to every given fine-grained cluster found during single-sample processing, and these average scores were further scaled (for plotting purposes) to the range 0– 1, by means of the following mapping:

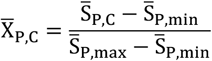

where 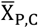 is the scaled (0–1) average score for a given pathway P in the fine-grained cluster C, while 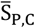 is the average pathway score for that cluster, and 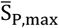 and 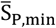 are respectively the highest and lowest values of 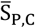 among all fine-grained clusters. For dynamic pathway-activity profiles, pathway activity scores were instead averaged over all cells in a given cell state, and these average scores were analogously scaled to the range 0–1 for plotting purposes.

### State-based and branch-based ranking of metabolic genes

The raw count matrix for the full merged data set was normalized to units of counts per 100k reads (CP100k), by dividing the UMI counts for every gene in each cell by the total UMI counts for that cell, and then multiplying times 10^5^. The latter was then subset to include only genes found within any of the metabolic gene sets in the KEGG metabolic pathway database (Kanehisa and Goto, 2000). The normalized expression values for those genes were then averaged over all cells annotated to a given cell state. A state-based score S_S,g_ was then determined for every metabolic gene g, based on how dominant the expression of that gene was in its most highly expressing cell state relative to all other states, using the formula

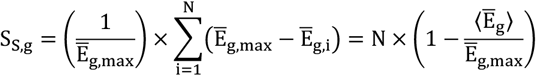

where i = 1, …, N denotes each of the different cell states, N is the total number of states considered (N = 11 here), Ē_g,i_ is the average normalized expression of gene g for cells in state i, Ē_g,max_ is the maximum value of Ē_g,i_ among all cell states (i.e. the value of Ē_g,i_ for the dominant state), and ⟨ Ē_g_⟩ is the average value of Ē_g,i_ over all cell states. Genes dominating in each state were then ranked by sorting them in descending order of their state-based score, to obtain ranked gene lists for every state. Similarly, branch-based scores S_B,g_ were determined for every metabolic gene based on the overall differences in relative expression between the differentiation branches leading to either a central-memory (CM = *Undiff + Tcm*_*p*_ *+ Tcm*) or an effector-memory (EM = *Teff + Tem*_*p*_ *+ Tem*) phenotype, using the formula

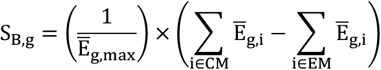

where each summation is restricted to cell states in either branch, and with positive/negative scores denoting respectively a dominant expression of gene g in the central/effector-memory branch. Genes dominating in each branch were then ranked by sorting them in descending order of the absolute value of their branch-based score, to obtain ranked gene lists for either branch. Genes with an average normalized expression below 1 count per 100k reads for all cell states (i.e. those with Ē_g,max_ < 1) were excluded from all rankings, to avoid any potential bias introduced by high relative oscillations in low expression genes.

### Gene expression profile correlations

The raw count matrix for the full merged data set was normalized to units of counts per 100k reads (CP100k) for each cell, by dividing the UMI counts for every gene in each cell by the total UMI counts for that cell, and then multiplying times 10^5^. Normalized expression values for those genes were then averaged over all cells annotated to a given cell state, to obtain expression profiles for all genes as a function of cell state. These expression profiles were further scaled to the range 0–1, by means of the following mapping:

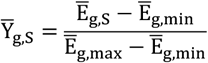

where 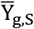 is the scaled (0–1) expression level for gene g in cell state S, while Ē_g,s_ is the average normalized expression level for that cluster, and Ē_g,max_ and Ē_g,min_ are respectively the highest and lowest values of Ē_g,s_ among all cell states. Pearson correlation coefficients and the corresponding p-values were then calculated in *R* between every pair of genes A and B of interest, using the function 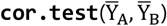 with 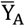 and 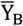 being vectors of the form 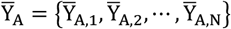 and analogously for 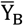.

### In vivo Asns gene-expression measurements

Naïve CD8^+^ T cells were isolated from the spleens of 6–12-week-old P14 transgenic mice using the MojoSort™ Mouse CD8 Naïve T Cell Isolation Kit, following the manufacturer’s guidelines, and 5×10^4^ cells were adoptively transferred into 5–8-week-old recipient C57BL/6J mice. 24 hours after transfer, mice were infected with 2×10^5^ PFU of LCMV Armstrong. At days 0, 5, 8, 15, and 28 post-infection, mice were sacrificed by cervical dislocation, and their spleens were harvested. CD8^+^ T cells were then isolated from the spleens using the MojoSort™ Mouse CD8 T Cell Isolation Kit (BioLegend), following the manufacturer’s guidelines, and immediately stained for FACS sorting and further RNA extraction, as described below.

### In vivo ASNS overexpression experiments

ASNS was stably overexpressed in activated CD8^+^ T cells using the murine stem cell virus (MSCV) retroviral expression system MSCV-IRES-Thy1.1 (kindly provided by Susan M. Kaech). Retroviral production was performed by transfection of Phoenix Eco cells with MSCV ASNS-overexpression (MSCV-ASNS-OE) or MSCV empty-vector (MSCV-EV, serving as control) constructs, with the aid of TurboFect™ Transfection Reagent (Thermo Scientific). Retroviral particles were collected 48 and 72h after transfection, concentrated by ultracentrifugation (2 hours at 24,000 x g, 4°C), and stored at -80°C until use.

Naïve CD8^+^ T cells were isolated from the spleens of 6–12-week-old P14 transgenic mice using the MojoSort™ Mouse CD8 Naïve T Cell Isolation Kit, following the manufacturer’s guidelines. Naïve CD8^+^ T cells were then activated *in vitro* using Dynabeads™ conjugated with anti-CD3 and anti-CD28 monoclonal antibodies (Gibco), for a total of 48h, in RPMI 1640 supplemented with 10% FBS, 100 U/mL penicillin, 100 µg/mL streptomycin, 1 mM sodium pyruvate, 2 mM L-glutamine, and 50 µM β-mercaptoethanol (all from Gibco). Concurrently with the start of *in vitro* activation, 5-week-old recipient C57BL/6J mice were infected with 2×10^5^ PFU of LCMV Armstrong. After 24h of stimulation, activated CD8^+^ T cells were transduced with MSCV-ASNS-OE or MSCV-EV viral particles on RetroNectin-coated plates (Takara Bio). 24 hours later, transduced cells were washed with PBS, and 5×10^4^ cells of each control and ASNS-OE were adoptively co-transferred into the time-matched infected recipient C57BL/6J mice. At days 8,12, and 28 post-infection, mice were sacrificed by cervical dislocation, and their spleens were immediately harvested and processed for FACS analysis, as described below.

### Flow cytometry

For *in vitro* activation/differentiation experiments, cells from the appropriate wells of the culture plates were transferred to a round-bottom 96-well plate (Corning 3799), washed once with PBS, resuspended in 50 µL/well of fixable Zombie Green viability dye (BioLegend) diluted 1/1000 in PBS, and stained for 15 minutes in the dark at room temperature. Cells were then washed twice with FACS buffer (PBS + 3% FBS + 2 mM EDTA), resuspended in 50 µL/well of surface-antibody cocktail, and stained for 30 minutes in the dark at 4°C. Surface-antibody cocktails were prepared in 2.4G2 hybridoma medium supplemented with 0.05% NaN_3_ (kindly provided by Stephanie-Humblet Baron), for simultaneous Fc-receptor blocking. Cells were then washed twice with FACS buffer, resuspended in a final volume of ∼100 µL/well of FACS buffer, and filter-transferred into 5 mL FACS tubes with cell-strainer caps (Falcon) for FACS analysis. Single-stained compensation controls were generated using 20 µL of UltraComp eBeads™ compensation beads (Invitrogen) per single-stain for all antibody-based stains, or a mixture of live and dead cells (the latter, achieved by warming up a small cell aliquot at 60°C for 15 minutes) for the fixable Zombie dye. Samples were acquired on a FACSCanto™ II analyzer (BD Biosciences) using the FACSDiva™ v8.0 software (BD Biosciences), and further analyzed with FlowJo v10.7 (BD Biosciences). Events were first gated to remove cell debris in the FSC-A *vs*. SSC-A plane, followed by selection of live (Zombie Green-negative) CD8^+^ T cells, removal of pro-apoptotic (low FSC) cells in the FSC-A *vs*. SSC-A plane, and doublet removal on the FSC-H *vs*. FSC-A plane. The following surface-marker antibodies (all from eBioscience) and dilutions were used for FACS analysis: anti-mouse CD8a APC (clone 53-6.7, 1/200 dilution), anti-mouse CD44 PE (clone IM7, 1/500 dilution), anti-mouse CD62L (L-Selectin) APC-eFluor 780 (clone MEL-14, 1/150 dilution), anti-mouse CD25 PerCP-Cyanine5.5 (clone PC61.5, 1/800 dilution), anti-mouse CD69 PE-Cyanine7 (clone H1.2F3, 1/200 dilution).

For *in vivo* ASNS overexpression experiments, splenocyte suspensions were prepared by mashing the freshly-resected whole spleens through a 180 µm iron-wire mesh (SEFAR) using the rubber ends of syringe plungers, followed by filtering through 70 µm nylon cell strainers. Cells were then resuspended in ACK red blood cell lysis buffer, incubated for 1 minute at room temperature, and then washed once and resuspended in FACS buffer. The resulting whole-splenocyte single-cell suspensions were directly stained in round-bottom 96-well plates using the following surface-marker antibodies (either from eBioscience, BioLegend, or produced in house): anti-mouse CD8a FITC (clone 53.6.7, 1/500 dilution), anti-mouse CD90.1 (Thy1.1) PE (clone HIS15, 1/500 dilution), anti-mouse CD45.1 Brilliant Violet 785™ (clone A20, 1/50 dilution), anti-mouse CD45.2 Pacific Blue (clone Ali 4A2, 1/100 dilution), anti-mouse CD44 APC-Cyanine7 (clone IM7, 1/100 dilution), anti-mouse CD62L (L-Selectin) Brilliant Violet 711™ (clone MEL-14, 1/1000 dilution), anti-mouse KLRG1 PE-Cyanine7 (clone 2F1, 1/500 dilution), anti-mouse CD127 Alexa Fluor^®^ 647 (clone A7R34, 1/100 dilution). Samples were then acquired on an LSR II™ analyzer (BD Biosciences) using the FACSDiva™ v8.0 software (BD Biosciences), and further analyzed with FlowJo v10.7 (BD Biosciences). Events were first gated in the FSC-A *vs*. SSC-A plane to select for lymphocytes and remove cell debris, followed by doublet removal on the FSC-H *vs*. FSC-W and SSC-H *vs*. SSC-W planes. Transduced CD8^+^ T cells were then selected as CD8/Thy1.1 double-positive cells, followed by gating on either ASNS-OE (CD45.1^Hi^CD45.2^Lo^) or control (CD45.1^Hi^ CD45.2^Hi^) cells.

For *in vivo Asns* gene-expression measurements, CD8^+^ T cells isolated at each time point of interest were directly stained in round-bottom 96-well plates using the following surface-marker antibodies (either from eBioscience, BioLegend, or produced in house): anti-mouse CD8a FITC (clone 53.6.7, 1/500 dilution), anti-mouse CD45.2 Pacific Blue (clone Ali 4A2, 1/100 dilution), anti-mouse CD45.1 PE (clone A20, 1/200 dilution), and either anti-mouse CD127 Alexa Fluor^®^ 647 (clone A7R34, 1/100 dilution) + anti-mouse KLRG1 PE-Cyanine7 (clone 2F1, 1/500 dilution), or anti-mouse CD44 APC (clone IM7, 1/500 dilution) + anti-mouse CD62L Pe-Cyanine7 (clone MEL-14, 1/1000 dilution). Cells were then sorted on a FACSAria III™ analyzer (BD Biosciences) using the FACSDiva™ v8.0 software (BD Biosciences). Events were first gated in the FSC-A *vs*. SSC-A plane to select for lymphocytes and remove cell debris, followed by doublet removal on the FSC-H *vs*. FSC-W and SSC-H *vs*. SSC-W planes. CD8^+^ T cells were then selected based on CD8 expression, followed by gating on adoptively-transferred cells (CD45.1^Hi^ CD45.2^Lo^). At day 8 post-infection, adoptively-transferred cells were further sorted based on KLRG1 and CD127 expression, to select for both short-lived effector cells (SLECs, KLRG1^Hi^ CD127^Lo^) and memory-precursor effector cells (MPECs, KLRG1^Lo^ CD127^Hi^). Similarly, at day 28 post-infection, adoptively-transferred cells were further sorted based on CD44 and CD62L expression, to select for both central-memory (Tcm, CD44^Hi^ CD62L^Hi^) and effector-memory (Tem, CD44^Hi^ CD62L^Lo^) cells. At days 0, 5, and 15 post-infection, the whole adoptively-transferred population was sorted without further subpopulation selection. Sorted cells were then lysed, and cell lysates were frozen until further used for RNA extraction, as described below.

### RNA extraction and qPCR

For *in vitro* activation/differentiation experiments, cell suspensions were pooled from multiple wells (∼0.3-1×10^6^ cells/replicate, depending on the amount of RNA expected per condition), and total RNA was extracted using the PureLink^®^ RNA Mini Kit (Invitrogen), following the manufacturer’s guidelines for ≤ 10^6^ cells/sample, including on-column PureLink^®^ DNase (Invitrogen) treatment, and with a final elution volume of 30 µL/sample. RNA concentration and purity were assessed based on a NanoDrop™ One (Thermo Scientific) spectrophotometer, after which RNA (∼300 ng/sample) was reverse-transcribed into cDNA using the qScript™ cDNA Synthesis Kit (Quantabio). Ct values for *Asns* and the housekeeping gene *Ppib* (Cyclophilin B) were determined by quantitative real-time PCR on a QuantStudio 12K Flex Real-Time PCR System (Applied Biosystems), using the Platinum^®^ SYBR^®^ Green qPCR SuperMix-UDG with ROX reference dye (Invitrogen) and specific primers (see **Supplementary Table 3**). Amplification was performed at 95°C for 2 minutes, followed by 40 cycles of 15 seconds at 95°C and 1 minute at 60°C. *Asns* expression levels were determined by first averaging the Ct values for *Asns* and *Ppib* over 3 technical replicates per biological replicate and condition, and then converting the resulting ΔCt (*Asns – Ppib*) values, for each biological replicate and condition, into housekeeper-normalized absolute expression levels X = 2^-ΔCt^. *Asns* expression levels (or fold-changes) relative to the naïve state were then determined by dividing the values of X for each biological replicate and condition by the average value of X for the naïve state.

For *in vivo Asns* gene-expression measurements, frozen cell lysates from the different cell subpopulations sorted at each time point were thawed, and RNA was isolated using the RNeasy^®^ Mini Kit (Qiagen), following the manufacturer’s guidelines, and with a final elution volume of 30 µL/sample. RNA concentration and purity were assessed based on a NanoDrop™ One (Thermo Scientific) spectrophotometer, and equal amounts of RNA for each sample were then reversed-transcribed using the PrimeScript™ RT Master Mix (Takara), following the manufacturer’s guidelines. Ct values for *Asns* and the housekeeping gene *Actb* (Beta-actin) were determined by quantitative real-time PCR on a LightCycler^®^ 480 II Real-Time PCR System (Roche Life Science), using the TB Green™ Premix Ex Taq™ (Takara) and specific primers (see **Supplementary Table 3**). Amplification was performed at 95°C for 30 seconds, followed by 40 cycles of 5 seconds at 95°C and 30 seconds at 60°C. *Asns* expression levels were determined by first averaging the Ct values for *Asns* and *Actb* over 2 technical replicates per biological replicate and condition, and then converting the resulting ΔCt (*Asns – Actb*) values, for each biological replicate and condition, into housekeeper-normalized absolute expression levels X = 2^-ΔCt^. *Asns* expression levels (or fold-changes) relative to the naïve state were then determined by dividing the values of X for each biological replicate and condition by the average value of X for the naïve state.

### Data presentation and statistical analysis

All data are presented as mean ± SD (standard deviation), mean ±SEM (standard error of the mean), or median + inter-quartile range (IQR), as indicated in the figure legends, with figures corresponding to *in vivo* data showing also individual data points. Statistical analysis was performed both within the *R/Bioconductor* framework, and using GraphPad Prism version 9 (GraphPad Software). The sample size for all experiments was chosen empirically, and at least 3 biological replicates (for *in vivo* data) or 3 independent culture wells (for *in vitro* data) were used for all statistical calculations. Unless noted, statistical analysis for all time-course profiles is based on linear regression of the data to orthogonal polynomials of the specified degree in the continuous time variable, either assuming joint coefficients for all conditions being compared (restricted model), or letting the coefficients depend on the condition (full model). Statistical comparisons between the full and restricted models are then performed using F-Tests, to determine significant differences between the time-course profiles for different conditions. This is done first considering all conditions together (global F-Test), and then between every pair of conditions (pairwise F-Tests), if more than 2, and provided that the global test was statistically significant (indicating a significant condition effect). Further statistical comparisons between data for different conditions at given (not necessarily fixed) time points are performed using multiple pairwise two-sided t-Tests with Welch’s correction (unequal variance assumption). Polynomial degrees for regression were chosen empirically for each data set to ensure goodness of fit, and were always set to at most 2 degrees lower than the number of time-points in the data set, to avoid overfitting. All p-values for pairwise F-Tests or t-Tests are subject to FDR-adjustment for multiple comparisons using the Benjamini-Hochberg method, considering all possible pairwise comparisons (even if reported for select comparisons only). Further details on the statistical tests applied to each data set are given in the corresponding figure legends. Statistical significance levels are indicated in the figures either by the corresponding p-values or using the following annotations: **ns**, not significant (p ≥ 0.05); **\***, p < 0.05; **\*\***, p < 0.01; **\*\*\***, p < 0.001.

## REFERENCES

Ahl, P.J., Hopkins, R.A., Xiang, W.W., Au, B., Kaliaperumal, N., Fairhurst, A.-M., and Connolly, J.E. (2020). Met-Flow, a Strategy for Single-Cell Metabolic Analysis Highlights Dynamic Changes in Immune Subpopulations. Commun. Biol. 3, 305.

Ahn, E., Araki, K., Hashimoto, M., Li, W., Riley, J.L., Cheung, J., Sharpe, A.H., Freeman, G.J., Irving, B.A., and Ahmed, R. (2018). Role of PD-1 During Effector CD8 T Cell Differentiation. Proc. Natl. Acad. Sci. 115, 4749–4754.

Aibar, S., González-Blas, C.B., Moerman, T., Huynh-Thu, V.A., Imrichova, H., Hulselmans, G., Rambow, F., Marine, J.-C., Geurts, P., Aerts, J., et al. (2017). SCENIC: Single-Cell Regulatory Network Inference and Clustering. Nat. Methods 14, 1083–1086.

Ananieva, E.A., Powell, J.D., and Hutson, S.M. (2016). Leucine Metabolism in T Cell Activation: mTOR Signaling and Beyond. Adv. Nutr. An Int. Rev. J. 7, 798S–805S.

Arruabarrena-Aristorena, A., Zabala-Letona, A., and Carracedo, A. (2018). Oil for the Cancer Engine: The Cross-Talk between Oncogenic Signaling and Polyamine Metabolism. Sci. Adv. 4, eaar2606.

Artyomov, M.N., and Van den Bossche, J. (2020). Immunometabolism in the Single-Cell Era. Cell Metab. 32, 710–725.

Bauer, D.E., Harris, M.H., Plas, D.R., Lum, J.J., Hammerman, P.S., Rathmell, J.C., Riley, J.L., and Thompson, C.B. (2004). Cytokine Stimulation of Aerobic Glycolysis in Hematopoietic Cells Exceeds Proliferative Demand. FASEB J. 18, 1303–1305.

Bergen, V., Lange, M., Peidli, S., Wolf, F.A., and Theis, F.J. (2020). Generalizing RNA Velocity to Transient Cell States through Dynamical Modeling. Nat. Biotechnol. 38, 1408–1414.

Best, J.A., Blair, D.A., Knell, J., Yang, E., Mayya, V., Doedens, A., Dustin, M.L., and Goldrath, A.W. (2013). Transcriptional Insights into the CD8+ T Cell Response to Infection and Memory T Cell Formation. Nat. Immunol. 14, 404–412.

Bjorkdahl, O., Barber, K.A., Brett, S.J., Daly, M.G., Plumpton, C., Elshourbagy, N.A., Tite, J.P., and Thomsen, L.L. (2003). Characterization of CC-Chemokine Receptor 7 Expression on Murine T Cells in Lymphoid Tissues. Immunology 110, 170–179.

Blondel, V.D., Guillaume, J.-L., Lambiotte, R., and Lefebvre, E. (2008). Fast Unfolding of Communities in Large Networks. J. Stat. Mech. Theory Exp. 2008, P10008.

Buck, M.D., O’Sullivan, D., and Pearce, E.L. (2015). T Cell Metabolism Drives Immunity. J. Exp. Med. 212, 1345–1360.

Butler, A., Hoffman, P., Smibert, P., Papalexi, E., and Satija, R. (2018). Integrating Single-Cell Transcriptomic Data across Different Conditions, Technologies, and Species. Nat. Biotechnol. 36, 411–420.

Cantor, J.R., Abu-Remaileh, M., Kanarek, N., Freinkman, E., Gao, X., Louissaint, A., Lewis, C.A., and Sabatini, D.M. (2017). Physiologic Medium Rewires Cellular Metabolism and Reveals Uric Acid as an Endogenous Inhibitor of UMP Synthase. Cell 169, 258-272.e17.

Cao, J., Spielmann, M., Qiu, X., Huang, X., Ibrahim, D.M., Hill, A.J., Zhang, F., Mundlos, S., Christiansen, L., Steemers, F.J., et al. (2019). The Single-Cell Transcriptional Landscape of Mammalian Organogenesis. Nature 566, 496–502.

Carr, E.L., Kelman, A., Wu, G.S., Gopaul, R., Senkevitch, E., Aghvanyan, A., Turay, A.M., and Frauwirth, K.A. (2010). Glutamine Uptake and Metabolism Are Coordinately Regulated by ERK/MAPK during T Lymphocyte Activation. J. Immunol. 185, 1037– 1044.

Cham, C.M., and Gajewski, T.F. (2005). Glucose Availability Regulates IFN-γ Production and p70S6 Kinase Activation in CD8+ Effector T Cells. J. Immunol. 174, 4670–4677.

Cham, C.M., Driessens, G., O’Keefe, J.P., and Gajewski, T.F. (2008). Glucose Deprivation Inhibits Multiple Key Gene Expression Events and Effector Functions in CD8+ T Cells. Eur. J. Immunol. 38, 2438–2450.

Chang, C.-H., Curtis, J.D., Maggi, L.B., Faubert, B., Villarino, A. V., O’Sullivan, D., Huang, S.C.-C., van der Windt, G.J.W., Blagih, J., Qiu, J., et al. (2013). Posttranscriptional Control of T Cell Effector Function by Aerobic Glycolysis. Cell 153, 1239–1251.

Chang, C.-H., Qiu, J., O’Sullivan, D., Buck, M.D., Noguchi, T., Curtis, J.D., Chen, Q., Gindin, M., Gubin, M.M., van der Windt, G.J.W., et al. (2015). Metabolic Competition in the Tumor Microenvironment Is a Driver of Cancer Progression. Cell 162, 1229–1241.

Chi, H. (2012). Regulation and Function of mTOR Signalling in T Cell Fate Decisions. Nat. Rev. Immunol. 12, 325–338.

Choi, B.-S., Martinez-Falero, I.C., Corset, C., Munder, M., Modolell, M., Muller, I., and Kropf, P. (2009). Differential Impact of L-Arginine Deprivation on the Activation and Effector Functions of T Cells and Macrophages. J. Leukoc. Biol. 85, 268–277.

Clarke, S.Rm., Barnden, M., Kurts, C., Carbone, F.R., Miller, J.F., and Heath, W.R. (2000). Characterization of the Ovalbumin-Specific TCR Transgenic Line OT-I: MHC Elements for Positive and Negative Selection. Immunol. Cell Biol. 78, 110–117.

Cornish, G.H., Sinclair, L. V., and Cantrell, D.A. (2006). Differential Regulation of T-Cell Growth by IL-2 and IL-15. Blood 108, 600–608.

Cui, W., and Kaech, S.M. (2010). Generation of Effector CD8+ T Cells and Their Conversion to Memory T Cells. Immunol. Rev. 236, 151–166.

Cui, G., Staron, M.M., Gray, S.M., Ho, P.-C., Amezquita, R.A., Wu, J., and Kaech, S.M. (2015). IL-7-Induced Glycerol Transport and TAG Synthesis Promotes Memory CD8+ T Cell Longevity. Cell 161, 750–761.

Edwards-Hicks, J., Mitterer, M., Pearce, E.L., and Buescher, J.M. (2020). Metabolic Dynamics of In Vitro CD8+ T Cell Activation. Metabolites 11, 12.

Elia, I., and Fendt, S.-M. (2016). In Vivo Cancer Metabolism is Defined by the Nutrient Microenvironment. Transl. Cancer Res. 5, S1284–S1287.

Fernández-García, J., and Fendt, S.-M. (2019). Assessing the impact of the nutrient microenvironment on the metabolism of effector CD8+ T cells.

Geiger, R., Rieckmann, J.C., Wolf, T., Basso, C., Feng, Y., Fuhrer, T., Kogadeeva, M., Picotti, P., Meissner, F., Mann, M., et al. (2016). L-Arginine Modulates T Cell Metabolism and Enhances Survival and Anti-tumor Activity. Cell 167, 829-842.e13.

Grün, D. (2018). Revealing Routes of Cellular Differentiation by Single-Cell RNA-seq. Curr. Opin. Syst. Biol. 11, 9–17.

Hänzelmann, S., Castelo, R., and Guinney, J. (2013). GSVA: Gene Set Variation Analysis for Microarray and RNA-Seq Data. BMC Bioinformatics 14, 7.

Hartmann, F.J., Mrdjen, D., McCaffrey, E., Glass, D.R., Greenwald, N.F., Bharadwaj, A., Khair, Z., Verberk, S.G.S., Baranski, A., Baskar, R., et al. (2021). Single-Cell Metabolic Profiling of Human Cytotoxic T Cells. Nat. Biotechnol. 39, 186–197.

Ho, P.-C., Bihuniak, J.D., Macintyre, A.N., Staron, M., Liu, X., Amezquita, R., Tsui, Y.-C., Cui, G., Micevic, G., Perales, J.C., et al. (2015). Phosphoenolpyruvate Is a Metabolic Checkpoint of Anti-tumor T Cell Responses. Cell 162, 1217–1228.

Hogquist, K.A., Jameson, S.C., Heath, W.R., Howard, J.L., Bevan, M.J., and Carbone, F.R. (1994). T Cell Receptor Antagonist Peptides Induce Positive Selection. Cell 76, 17–27.

Hope, H.C., Brownlie, R.J., Steele, L., and Salmond, R.J. (2020). Coordination of Asparagine Uptake and Asparagine Synthetase Expression is Required for T Cell Activation. BioRxiv.

Hope, H.C., Brownlie, R.J., Fife, C.M., Steele, L., Lorger, M., and Salmond, R.J. (2021). Coordination of Asparagine Uptake and Asparagine Synthetase Expression Modulates CD8+ T Cell Activation. JCI Insight.

Jones, N., Cronin, J.G., Dolton, G., Panetti, S., Schauenburg, A.J., Galloway, S.A.E., Sewell, A.K., Cole, D.K., Thornton, C.A., and Francis, N.J. (2017). Metabolic Adaptation of Human CD4+ and CD8+ T-Cells to T-Cell Receptor-Mediated Stimulation. Front. Immunol. 8, 1516.

Kaech, S.M., and Cui, W. (2012). Transcriptional Control of Effector and Memory CD8+ T Cell Differentiation. Nat. Rev. Immunol. 12, 749–761.

Kakaradov, B., Arsenio, J., Widjaja, C.E., He, Z., Aigner, S., Metz, P.J., Yu, B., Wehrens, E.J., Lopez, J., Kim, S.H., et al. (2017). Early Transcriptional and Epigenetic Regulation of CD8+ T Cell Differentiation Revealed by Single-Cell RNA Sequencing. Nat. Immunol. 18, 422–432.

Kalia, V., Sarkar, S., Subramaniam, S., Haining, W.N., Smith, K.A., and Ahmed, R. (2010). Prolonged Interleukin-2Rα Expression on Virus-Specific CD8+ T Cells Favors Terminal-Effector Differentiation In Vivo. Immunity 32, 91–103.

Kanehisa, M., and Goto, S. (2000). KEGG: Kyoto Encyclopedia of Genes and Genomes. Nucleic Acids Res. 28, 27–30.

Kelly, B., and Pearce, E.L. (2020). Amino Assets: How Amino Acids Support Immunity. Cell Metab. 32, 154–175.

Kharchenko, P. V., Silberstein, L., and Scadden, D.T. (2014). Bayesian Approach to Single-Cell Differential Expression Analysis. Nat. Methods 11, 740–742.

Kidani, Y., Elsaesser, H., Hock, M.B., Vergnes, L., Williams, K.J., Argus, J.P., Marbois, B.N., Komisopoulou, E., Wilson, E.B., Osborne, T.F., et al. (2013). Sterol Regulatory Element–Binding Proteins are Essential for the Metabolic Programming of Effector T Cells and Adaptive Immunity. Nat. Immunol. 14, 489–499.

Kim, T.K., and Eberwine, J.H. (2010). Mammalian Cell Transfection: The Present and the Future. Anal. Bioanal. Chem. 397, 3173–3178.

Krall, A.S., Xu, S., Graeber, T.G., Braas, D., and Christofk, H.R. (2016). Asparagine Promotes Cancer Cell Proliferation through Use as an Amino Acid Exchange Factor. Nat. Commun. 7, 11457.

Krall, A.S., Mullen, P.J., Surjono, F., Momcilovic, M., Schmid, E.W., Halbrook, C.J., Thambundit, A., Mittelman, S.D., Lyssiotis, C.A., Shackelford, D.B., et al. (2021). Asparagine Couples Mitochondrial Respiration to ATF4 Activity and Tumor Growth. Cell Metab. 33, 1013-1026.e6.

Leney-Greene, M.A., Boddapati, A.K., Su, H.C., Cantor, J.R., and Lenardo, M.J. (2020). Human Plasma-like Medium Improves T Lymphocyte Activation. IScience 23, 100759.

Levine, L.S., Hiam-Galvez, K.J., Marquez, D.M., Tenvooren, I., Madden, M.Z., Contreras, D.C., Dahunsi, D.O., Irish, J.M., Oluwole, O.O., Rathmell, J.C., et al. (2021). Single-Cell Analysis by Mass Cytometry Reveals Metabolic States of Early-Activated CD8+ T Cells During the Primary Immune Response. Immunity 54, 829-844.e5.

Liu, Q., Sun, Z., and Chen, L. (2020). Memory T Cells: Strategies for Optimizing Tumor Immunotherapy. Protein Cell 11, 549– 564.

Loftus, R.M., and Finlay, D.K. (2016). Immunometabolism: Cellular Metabolism Turns Immune Regulator. J. Biol. Chem. 291, 1–10.

Ma, E.H., Bantug, G., Griss, T., Condotta, S., Johnson, R.M., Samborska, B., Mainolfi, N., Suri, V., Guak, H., Balmer, M.L., et al. (2017). Serine Is an Essential Metabolite for Effector T Cell Expansion. Cell Metab. 25, 1–13.

Ma, E.H., Verway, M.J., Johnson, R.M., Roy, D.G., Steadman, M., Hayes, S., Williams, K.S., Sheldon, R.D., Samborska, B., Kosinski, P.A., et al. (2019). Metabolic Profiling Using Stable Isotope Tracing Reveals Distinct Patterns of Glucose Utilization by Physiologically Activated CD8+ T Cells. Immunity 51, 856-870.e5.

Ma, R., Ji, T., Zhang, H., Dong, W., Chen, X., Xu, P., Chen, D., Liang, X., Yin, X., Liu, Y., et al. (2018). A Pck1-Directed Glycogen Metabolic Program Regulates Formation and Maintenance of Memory CD8+ T Cells. Nat. Cell Biol. 20, 21–27.

MacIver, N.J., Michalek, R.D., and Rathmell, J.C. (2013). Metabolic Regulation of T Lymphocytes. Annu. Rev. Immunol. 31, 259–283.

La Manno, G., Soldatov, R., Zeisel, A., Braun, E., Hochgerner, H., Petukhov, V., Lidschreiber, K., Kastriti, M.E., Lönnerberg, P., Furlan, A., et al. (2018). RNA Velocity of Single Cells. Nature 560, 494–498.

Marguerat, S., Schmidt, A., Codlin, S., Chen, W., Aebersold, R., and Bähler, J. (2012). Quantitative Analysis of Fission Yeast Transcriptomes and Proteomes in Proliferating and Quiescent Cells. Cell 151, 671–683.

Marzo, A.L., Klonowski, K.D., Bon, A. Le, Borrow, P., Tough, D.F., and Lefrançois, L. (2005). Initial T Cell Frequency Dictates Memory CD8+ T Cell Lineage Commitment. Nat. Immunol. 6, 793–799.

McInnes, L., Healy, J., and Melville, J. (2018). UMAP: Uniform Manifold Approximation and Projection for Dimension Reduction. ArXiv.

Meng, D., Yang, Q., Wang, H., Melick, C.H., Navlani, R., Frank, A.R., and Jewell, J.L. (2020). Glutamine and Asparagine activate mTORC1 Independently of Rag GTPases. J. Biol. Chem. 295, 2890–2899.

Mombaerts, P., Clarke, A.R., Rudnicki, M.A., Iacomini, J., Itohara, S., Lafaille, J.J., Wang, L., Ichikawa, Y., Jaenisch, R., Hooper, M.L., et al. (1992). Mutations in T-Cell Antigen Receptor Genes α and β Block Thymocyte Development at Different Stages. Nature 360, 225–231.

Monterisi, S., Lobo, M.J., Livie, C., Castle, J.C., Weinberger, M., Baillie, G., Surdo, N.C., Musheshe, N., Stangherlin, A., Gottlieb, E., et al. (2017). PDE2A2 Regulates mitochondria Morphology and Apoptotic Cell Death via Local Modulation of cAMP/PKA Signalling. Elife 6, 1–20.

Mueller, K., Schweier, O., and Pircher, H. (2008). Efficacy of IL-2-versus IL-15-Stimulated CD8 T Cells in Adoptive Immunotherapy. Eur. J. Immunol. 38, 2874–2885.

O’Sullivan, D. (2019). The Metabolic Spectrum of Memory T Cells. Immunol. Cell Biol. 97, 636–646.

O’Sullivan, D., van der Windt, G.J.W., Huang, S.C.-C., Curtis, J.D., Chang, C.-H., Buck, M.D., Qiu, J., Smith, A.M., Lam, W.Y., DiPlato, L.M., et al. (2014). Memory CD8+ T Cells Use Cell-Intrinsic Lipolysis to Support the Metabolic Programming Necessary for Development. Immunity 41, 75–88.

Obar, J.J., and Lefrançois, L. (2010). Memory CD8+ T Cell Differentiation. Ann. N. Y. Acad. Sci. 1183, 251–266.

Parham, P. (2014). The Immune System (New York: Garland Science).

Pearce, E.L. (2010). Metabolism in T Cell Activation and Differentiation. Curr. Opin. Immunol. 22, 314–320.

Pearce, E.J., and Pearce, E.L. (2017). Driving Immunity: All Roads Lead to Metabolism. Nat. Rev. Immunol. 18, 81–82.

Pearce, E.L., and Pearce, E.J. (2013). Metabolic pathways in immune cell activation and quiescence. Immunity 38, 633–643.

Pearce, E.L., Walsh, M.C., Cejas, P.J., Harms, G.M., Shen, H., Wang, L.-S., Jones, R.G., and Choi, Y. (2009). Enhancing CD8 T-cell Memory by Modulating Fatty Acid Metabolism. Nature 460, 103–107.

Pearce, E.L., Poffenberger, M.C., Chang, C.-H., and Jones, R.G. (2013). Fueling Immunity: Insights into Metabolism and Lymphocyte Function. Science (80-.). 342, 1242454.

Pegg, A.E. (2016). Functions of Polyamines in Mammals. J. Biol. Chem. 291, 14904–14912.

Pircher, H., Bürki, K., Lang, R., Hengartner, H., and Zinkernagel, R.M. (1989). Tolerance Induction in Double Specific T-Cell Receptor Transgenic Mice Varies with Antigen. Nature 342, 559–561.

Puleston, D.J., Villa, M., and Pearce, E.L. (2017). Ancillary Activity: Beyond Core Metabolism in Immune Cells. Cell Metab. 26, 131–141.

Rao, R.R., Li, Q., Odunsi, K., and Shrikant, P.A. (2010). The mTOR Kinase Determines Effector versus Memory CD8 + T Cell Fate by Regulating the Expression of Transcription Factors T-bet and Eomesodermin. Immunity 32, 67–78.

Raynor, J.L., and Chi, H. (2021). LCK Senses Asparagine for T Cell Activation. Nat. Cell Biol. 23, 7–8.

Rinaldi, G., Rossi, M., and Fendt, S.-M. (2018). Metabolic Interactions in Cancer: Cellular Metabolism at the Interface Between the Microenvironment, the Cancer Cell Phenotype and the Epigenetic Landscape. Wiley Interdiscip. Rev. Syst. Biol. Med. 10, e1397.

Ron-Harel, N., Ghergurovich, J.M., Notarangelo, G., LaFleur, M.W., Tsubosaka, Y., Sharpe, A.H., Rabinowitz, J.D., and Haigis, M.C. (2019). T Cell Activation Depends on Extracellular Alanine. Cell Rep. 28, 3011-3021.e4.

Rossi, A., Pacella, I., and Piconese, S. (2021). RNA Flow Cytometry for the Study of T Cell Metabolism. Int. J. Mol. Sci. 22, 3906.

Sinclair, L. V., Howden, A.J.M., Brenes, A., Spinelli, L., Hukelmann, J.L., Macintyre, A.N., Liu, X., Thomson, S., Taylor, P.M., Rathmell, J.C., et al. (2019). Antigen Receptor Control of Methionine Metabolism in T Cells. Elife 8, 1–29.

Stubbington, M.J.T., Rozenblatt-Rosen, O., Regev, A., and Teichmann, S.A. (2017). Single-Cell Transcriptomics to Explore the Immune System in Health and Disease. Science (80-.). 358, 58–63.

Szabo, P.A., Levitin, H.M., Miron, M., Snyder, M.E., Senda, T., Yuan, J., Cheng, Y.L., Bush, E.C., Dogra, P., Thapa, P., et al. (2019). Single-Cell Transcriptomics of Human T Cells Reveals Tissue and Activation Signatures in Health and Disease. Nat. Commun. 10, 4706.

Tardito, S., Oudin, A., Ahmed, S.U., Fack, F., Keunen, O., Zheng, L., Miletic, H., Sakariassen, P.Ø., Weinstock, A., Wagner, A., et al. (2015). Glutamine Synthetase Activity Fuels Nucleotide Biosynthesis and Supports Growth of Glutamine-Restricted Glioblastoma. Nat. Cell Biol. 17, 1556–1568.

Thurnher, M., and Gruenbacher, G. (2015). T Lymphocyte Regulation by Mevalonate Metabolism. Sci. Signal. 8, 1–10.

Torres, A., Luke, J.D., Kullas, A.L., Kapilashrami, K., Botbol, Y., Koller, A., Tonge, P.J., Chen, E.I., Macian, F., and van der Velden, A.W.M. (2016). Asparagine Deprivation Mediated by Salmonella Asparaginase Causes Suppression of Activation-Induced T Cell Metabolic Reprogramming. J. Leukoc. Biol. 99, 387–398.

Veiga-Fernandes, H., Walter, U., Bourgeois, C., McLean, A., and Rocha, B. (2000). Response of Naïve and Memory CD8+ T Cells to Antigen Stimulation In Vivo. Nat. Immunol. 1, 47–53.

Vande Voorde, J., Ackermann, T., Pfetzer, N., Sumpton, D., Mackay, G., Kalna, G., Nixon, C., Blyth, K., Gottlieb, E., and Tardito, S. (2019). Improving the Metabolic Fidelity of Cancer Models with a Physiological Cell Culture Medium. Sci. Adv. 5, eaau7314.

Waickman, A.T., and Powell, J.D. (2012). mTOR, Metabolism, and the Regulation of T-cell Differentiation and Function. Immunol. Rev. 249, 43–58.

Wang, R., and Green, D.R. (2012). Metabolic Reprogramming and Metabolic Dependency in T Cells. Immunol. Rev. 249, 14– 26.

Wang, R., Dillon, C.P., Shi, L.Z., Milasta, S., Carter, R., Finkelstein, D., McCormick, L.L., Fitzgerald, P., Chi, H., Munger, J., et al. (2011). The Transcription Factor Myc Controls Metabolic Reprogramming upon T Lymphocyte Activation. Immunity 35, 871– 882.

Wei, J., Raynor, J., Nguyen, T.-L.M., and Chi, H. (2017). Nutrient and Metabolic Sensing in T Cell Responses. Front. Immunol. 8, 1–14.

Weninger, W., Crowley, M.A., Manjunath, N., and von Andrian, U.H. (2001). Migratory Properties of Naive, Effector, and Memory CD8+ T Cells. J. Exp. Med. 194, 953–966.

Wherry, E.J., Teichgräber, V., Becker, T.C., Masopust, D., Kaech, S.M., Antia, R., von Andrian, U.H., and Ahmed, R. (2003). Lineage Relationship and Protective Immunity of Memory CD8 T Cell Subsets. Nat. Immunol. 4, 225–234.

van der Windt, G.J.W., and Pearce, E.L. (2012). Metabolic switching and fuel choice during T-cell differentiation and memory development. Immunol. Rev. 249, 27–42.

van der Windt, G.J.W., Everts, B., Chang, C.-H., Curtis, J.D., Freitas, T.C., Amiel, E., Pearce, E.J., and Pearce, E.L. (2012). Mitochondrial Respiratory Capacity Is a Critical Regulator of CD8+ T Cell Memory Development. Immunity 36, 68–78.

Wu, J., Li, G., Li, L., Li, D., Dong, Z., and Jiang, P. (2021). Asparagine Enhances LCK Signalling to Potentiate CD8+ T-Cell Activation and Anti-Tumour Responses. Nat. Cell Biol. 23, 75–86.

Zhang, H., Tang, K., Ma, J., Zhou, L., Liu, J., Zeng, L., Zhu, L., Xu, P., Chen, J., Wei, K., et al. (2020). Ketogenesis-Generated β-Hydroxybutyrate is an Epigenetic Regulator of CD8+ T-Cell Memory Development. Nat. Cell Biol. 22, 18–25.

Zhang, X., Sun, S., Hwang, I., Tough, D.F., and Sprent, J. (1998). Potent and Selective Stimulation of Memory-Phenotype CD8+ T Cells In Vivo by IL-15. Immunity 8, 591–599.

